# Receptor kinase LecRK-I.9 regulates cell wall remodelling and signalling during lateral root formation in *Arabidopsis*

**DOI:** 10.1101/2024.01.31.578125

**Authors:** Kevin Bellande, David Roujol, Josiane Chourré, Sophie Le Gall, Yves Martinez, Alain Jauneau, Vincent Burlat, Elisabeth Jamet, Hervé Canut

## Abstract

Assembling and remodelling the cell wall is essential for plant development. Cell wall dynamic is controlled by cell wall proteins and a variety of sensor and receptor systems. LecRK-I.9, an *Arabidopsis thaliana* plasma membrane-localised lectin receptor kinase, was previously shown to be involved in cell wall-plasma membrane contacts and to play roles in plant-pathogen interactions, but so far, its role in development was unknown. *LecRK-I.9* is transcribed at a high level in root tissues including the pericycle. Comparative transcript profiling of a loss-of-function mutant *vs* wild type identifies LecRK-I.9 as a regulator of cell wall metabolism. Consistently, *lecrk-I.9* mutants display an increased pectin methylesterification level correlated with decreased pectin methylesterase and increased polygalacturonase activities. Also, LecRK-I.9 impacts lateral root development through the regulation of genes encoding (*i*) cell wall remodelling proteins during early events of lateral root initiation, and (*ii*) cell wall signalling peptides (CLE2, CLE4) repressing lateral root emergence and growth. Besides, low nitrate reduces *LecRK-I.9* expression in pericycle and interferes with its regulatory network: however, the control of *CLE2* and *CLE4* expression is maintained. Altogether, the results show that LecRK-I.9 is a key player in a signalling network regulating both pre-branch site formation and lateral root emergence.

**Highlight:** The lectin receptor kinase LecRK-I.9 regulates the molecular events leading to lateral root formation in both the initiation and emergence processes in Arabidopsis through cell wall remodelling enzymes and signalling peptides.

## Introduction

As cells grow and differentiate, CWs are modified in their biochemical composition and structure to ensure cell functions, *i.e.* maintain cell integrity, allow cell-to-cell communication, adjust cell-to-cell adhesion, and support cell division and growth. Hence, CW dynamics largely control tissue morphology and plant development (Wolf *et al*., 2012).

Current models for the CW structure of growing cells consist of a highly hydrated matrix based on a scaffold of cellulose microfibrils that are cross-linked by branched hemicelluloses, such as xyloglucans (XGs), and embedded in pectins, such as homogalacturonan (HG) and rhamnogalacturonans I and II (RG-I and RG-II). Crosslinks between CW polysaccharides are either non-covalent (*e.g.* cellulose chains hydrogen-bonded together to form microfibrils, XGs hydrogen-bond to microfibrils, HG cross-linked by Ca^2+^ bridges) or covalent (*e.g.* pairs of RG-II by borate-diester bonds) (Fry, 2004; Sanhueza *et al*., 2022). It has also been proposed that the XGs-cellulose interactions mainly occur at the level of biomechanical hot spots (Cosgrove, 2016). The strength of these interactions determines the mechanical properties of the CW, often referred to as CW extensibility, and its capacity to balance the force exerted by the turgor pressure, which is the motive force for growth (Wolf *et al*., 2012). Interestingly, it was recently shown that the CW contains pectin nanofilaments that possess an intrinsic expansion capacity able to drive morphogenesis without turgor pressure (Haas *et al*., 2020). Due to the complexity of the CW architecture, CW dynamics relies on many parameters such as the deposition of new CW components (Lampugnani *et al*., 2018), the remodelling of the existing structures which is achieved through the activity of various CW remodelling proteins (Cosgrove, 2005), the hydration status (Wolf *et al*., 2012), or the availability of calcium and boron ions (Lamport *et al*., 2018; Voxeur & Fry, 2014). A strict coordination of these processes is required, including biosynthesis and secretion of the polysaccharides and/or the remodelling enzymes, as well as communication between cells. Then, CW dynamics must be tightly regulated for proper morphogenesis at cellular and tissue levels (Boudon *et al*., 2015).

The number of lateral roots (LRs) is considered a major agronomic trait because it directly influences crop yield (Zhu *et al*., 2011). In *Arabidopsis thaliana*, LRs grow from competent xylem-pole pericycle (XPP) cells, a single-cell layer that resides deep within the primary root. LR development follows a sequence of five main steps, (1) pre-branch site formation, (2) initiation, (3) morphogenesis, (4) emergence and (5) growth (Banda *et al*., 2019). LRs grow through the overlying endodermal, cortical and epidermal cell layers thus requiring extensive CW remodelling (Vermeer *et al*., 2014). Genes encoding CW remodelling proteins are locally induced at the earliest (Wachsman *et al*., 2020; Ramakrishna *et al*., 2019) and later steps (Swarup *et al*., 2008). In particular, genes encoding CW enzymes acting on HG are spatially and temporarily induced at the pre-branch sites: these enzymes are probably responsible for the differential distribution of esterified and de-esterified pectins at the sites of LR primordia (LRP) initiation (Wachsman *et al*., 2020). It is assumed that such a differential distribution could reduce cell adhesion of overlaying tissues, thus allowing LR emergence and, in contrast, stiffening the opposite tissues to avoid any disruption. The CW structures also need to be adapted during the swelling of the founder cells and the shrinking of the adjacent endodermal cells that are required to form LRPs (Vermeer *et al*., 2014; Ramakrishna *et al*., 2019).

LR development can only be achieved by the interconnection of communication networks. Until recently, much of the focus has been put on the phytohormone auxin and reactive oxygen species (ROS), but accumulated evidence in *A. thaliana* indicates that pairs of CW signalling peptides and their receptors are also involved in the control of LR development (Jourquin *et al*., 2020; Ou *et al*., 2021). At each developmental phase, auxin plays an important role (Du & Scheres, 2018). For instance, an auxin source originating from the primary root cap cells stimulates the formation of the LR pre-branch sites in XPP cells (Xuan *et al*., 2015). Besides, ROS have a strong impact on CW properties. They can be produced by RESPIRATORY BURST OXIDASE HOMOLOG (RBOH) plasma membrane proteins and CW class III peroxidases (CIII Prxs) through the dual hydroxylic and peroxidative cycles leading to either CW loosening or CW stiffening (Francoz *et al*., 2015). Both *RBOH* and *CIII Prx* genes appear to regulate ROS production in the overlaying tissue to facilitate LR emergence (Manzano *et al*., 2014; Orman-Ligeza *et al*., 2016).

THESEUS 1 (THE1), a plasma membrane receptor-like kinase (RLK) with an extracellular malectin domain, and its ligand RAPID ALKALINIZATION FACTOR 34 (RALF 34) regulate LR initiation by fine-tuning the asymmetric divisions of the founder cells (Murphy *et al*., 2016; Gonneau *et al*., 2018). The signalling module RALF34/THE1 also seems to depend on FERONIA (FER) another RLK with an extracellular malectin domain that is known to perceive other RALF peptides (Gonneau *et al*., 2018). Together with FER, THE1 is thought to be part of a surveillance system of the CW (Wolf & Höfte, 2014). A second signalling module example is provided by the INFLORESCENCE DEFICIENT IN ABSCISSION (IDA) CW peptide and its receptors HAESA (HAE) and HAESA-LIKE2 (HSL2), both belonging to the leucine-reach repeat (LRR)-RLK family. This module can promote cell separation in the tissues overlaying the LR primordium by up-regulating genes encoding CW remodelling proteins (Kumpf *et al*., 2013). Indeed, *IDA*, *HAE* and *HSL2* are expressed in the tissue layers covering the LR primordium and are up-regulated by auxin (Kumpf *et al*., 2013). A third signalling module composed of CLAVATA3/ESR-RELATED PROTEINS (CLEs) and CLAVATA1 (CLV1) LRR-RLK acts as a negative regulator of LR primordium growth and elongation without affecting LR initiation (Araya *et al*., 2014). *CLE2*, *4*, *5*, *6* and *7* are repressed upon nitrogen deficiency (Ma *et al*., 2020) suggesting that the CLE/CLV1 module links the perception of environmental cues and LR development. Despite these notable examples, however, a comprehensive list of the receptors involved in CW changes during plant development is still missing.

LecRK-I.9, a Legume-type lectin receptor kinase of *A. thaliana*, has previously been shown to mediate CW-plasma membrane contacts through protein-protein interactions suggesting both a structural and a signalling role at the cell surface (Gouget *et al*., 2006). LecRK-I.9 also plays important roles in plant innate immunity suggesting that the proteins maintaining CW-plasma membrane contacts also function in plant defence (Bouwmeester *et al*., 2011; Balagué *et al*., 2017). While the ability of LecRK-I.9 to bind complex carbohydrates is assumed, but not experimentally proven (Gouget *et al*., 2006), LecRK-I.9 was identified as a receptor for extracellular ATP and alternatively named DORN1 (Choi *et al*., 2014). It is assumed that ATP is released during physical damage of cells as a danger signal.

This study shows that LecRK-I.9 also controls LR development. Null mutants for *LecRK-I.9* show alterations in the number of LRs and in their density, as well as in the methylesterification of HG and the activities of CW remodelling enzymes. A transcript abundance analysis, comparing a *lecrk-I.9-1* mutant and wild-type plants, revealed the de-regulation of about 450 genes, one-third of which encoding CW-related proteins. Of particular interest was a group of up-regulated genes in the mutant that encode CW remodelling enzymes, but also a group of down-regulated genes that encode signalling peptides among which *CLE2* and *CLE4*. We conclude that LecRK-I.9 is part of a signalling network negatively regulating both pre-branch site formation and LR emergence and growth.

## Materials and methods

### Plant materials and growth conditions

All the experiments were performed using the *A. thaliana* ecotype Col-0 plants. Seeds of the T-DNA insertion lines *lecrk-I.9-1* (SALK_042209) and *lecrk-I.9-2* (SALK_024581) (Balagué *et al*., 2017) were obtained from the Nottingham Arabidopsis Stock Centre. The *CaMV35S:: LecRK-I.9* overexpressor (*Ox-1*; *Ox-2*) and the *proLecRK-I.9::GUS* lines were previously described (Bouwmeester *et al*., 2011; Balagué *et al*., 2017).

Seeds (15 mg) were surface-sterilized with 1 mL 2% sodium hypochlorite for 20 min. After five washes with sterile tap water, the seeds were placed in 250 mL Erlenmeyer flasks containing 50 mL of a half Murashige and Skoog (MS) medium (20 mM KNO_3_) (Sigma-Aldrich M5519, St Louis, MO, USA) supplemented with 1% sucrose (w/v) at pH 5.7 (KOH). The seeds were germinated and grown in flasks placed on a rotary shaker (130 rpm, New Brunswick orbital shaker model G 10-21, Edison, NJ, USA) under a photoperiod of 16 h (75 μmol.m^-2^ .s^-1^)/8 h dark at 20°C for 7 days. Low-concentrated nitrate media were made using half nitrogen-free MS salts (Sigma-Aldrich M0529) supplemented with 1.5 mM CaCl_2_, 0.75 mM MgSO_4_, 0.5 mM KH_2_PO_4_, 10 mM KCl, 0.5 mL/L organics (Sigma-Aldrich M3900), either 1 mM or 0.1 mM KNO_3_ and 1% sucrose (w/v) at pH 5.7 (KOH).

To obtain adventitious roots, seeds were sown on 1.2% (w/v) agar plates containing ½ MS medium supplemented with 1% sucrose (w/v) at pH 5.7 (KOH). The seeds were germinated and grown in the dark for 4 days. Etiolated seedlings were then transferred to new agar plates containing an MS-based medium (see above) with 1 mM KNO_3_. Etiolated seedlings were lying flat on the solid medium and the plates were placed vertically under a photoperiod of 16 h (75 μmol.m^−2^.s^−1^)/8 h dark at 25°C for 7 days.

### Generation of constructs

All the constructs were generated using the gateway® cloning technology.

To make the *proLecRK-I.9::LecRK-I.9::tagRFP* construct (*LecRK-I.9*: *At5g60300*), a 1486 bp genomic DNA fragment upstream of the start codon of *LecRK-I.9* together with its coding sequence was amplified by PCR. The same DNA fragment was used to complement *lecrk-I.9-1* and *lecrk-I.9-2*.

To make the *proLecRK-I.9::LecRK-I.9*Δ*kinase::tagRFP* construct, a 1486 bp genomic DNA fragment upstream of the start codon of *LecRK-I.9* together with part of the coding sequence of *LecRK-I.*9 including the signal peptide, the Legume lectin domain, the transmembrane domain and the arrest sequence, but omitting the kinase domain and the C-term extension was amplified by PCR. This construct was used to create dominant negative mutant lines.

To create the *proPME2::eGFP::GUS* reporter (*PME2*: *At1g53830*), the *proCLE2::eGFP::GUS* reporter (*CLE2*: *At4g18510*) and the *proCLE4::eGFP::GUS* reporter (*CLE4*: *At2g31081*), DNA fragments carrying 1562 bp, 1546 bp and 1556 bp genomic DNA upstream of their respective start codons were amplified by PCR, respectively.

All the fragments were amplified by PCR from the *A. thaliana* Col-0 genomic DNA using the AccuPrime™ *Taq* DNA Polymerase High Fidelity (Invitrogen®, Carlsbad, CA, USA). The primer sequences are given in Supplementary Table S1. All the PCR fragments were cloned in the pDONR207 entry vector (Invitrogen®), except for the PCR fragments to make the *proLecRK-I.9::LecRK-I.9::tagRFP* and the *proLecRK-I.9::LecRK-I.9*Δ*kinase::tagRFP* constructs which were inserted between the *EcoR*I and *BamH*I sites of the Gateway® tagRFP-AS-N entry vector (Evrogen, Moscow, Russia). After each transformation of *Escherichia coli*, the plasmids obtained from single colonies were sequenced. Expression vectors were generated by LR reactions using the pDONR207 entry clones and the destination vector pKGWFS7 (no promoter, *eGFP* and *GUS* reporters) (www.psb.ugent.be/Gateway) or the tagRFP-AS-N entry clones and the binary vector pAM-PAT-D35S-GWY-3HA-RTerm (gift from L. Deslandes, INRAe, Auzeville-Tolosane, France). The *Agrobacterium tumefaciens* GV3101::pMP90 strain was transformed with the recombinant vectors and used to stably transform WT, *lecrk-I.9-1* or *lecrk-I.9-2* plants with the floral dip method. T2 and T3 progenies were used for analysis.

The *DR5::GUS/lecRK-I.9-1*, *DR5::GUS/lecRK-I.9-2 and DR5::GUS/lecRK-I.9-2*Δ*kinase* plants were obtained by crossing experiments using *DR5::GUS* plants (Ulmasov *et al.,* 1997).

### Microarray analysis and data processing

Col-0 and *lecrk-I.9-1* seedlings were grown from seeds for 7 days in 250 mL Erlenmeyer flasks containing 50 mL of a half Murashige and Skoog (MS) medium (20 mM KNO_3_) (Sigma-Aldrich M5519, St Louis, MO, USA) supplemented with 1% sucrose (w/v) at pH 5.7 (KOH). Three biological replicates were conducted with one flask yielding a single biological replicate. Upon harvest, plant material was washed with UHQ water. For each flask, ∼1000 root segments between the root tip and the root-hypocotyl junction were pooled, frozen in liquid nitrogen, and stored at −80°C until use.

The Col-0 and *lecrk-I.9-1* root samples were sent to OakLabs (Hennigsdorf, Germany), where total RNA isolation, sample preparation and hybridization to ArrayXS Arabidopsis were performed on January 2016 (http://www.oak-labs.com/) (Supplementary Dataset 1). ArrayXS Arabidopsis is an 8x60K Agilent microarray which contains 32072 gene-specific tags, corresponding to 30541 gene loci.

### RNA extraction and quantitative RT-PCR

Total RNA was extracted from roots sampled as above using the SV total RNA isolation kit system following the manufacturer’s instructions (Promega, Madison, WI, USA). cDNA was synthesized with Superscript III reverse transcriptase (Invitrogen®) using 1 µg of total RNA. Real-time quantitative PCR (qPCR) was performed on a QuantStudio6 apparatus (Applied Biosystems®, Carlsbad, CA, USA) using Takyon™ reagents (Eurogentec, Liège, Belgium). MasterMix, 5 pmol of each primer and 1 µL of a 5-fold dilution of RT reaction product were in a 10 µL final reaction volume with PCR cycling conditions as follows: 10 min at 95°C, followed by 40 cycles of 15 s at 95°C, 60 s at 60°C. The primers are listed in Supplementary Table S1. All the reactions were checked for their dissociation curves. ΔCT between each gene and the internal controls (*At2g28390*, *At4g33380* and *At5g55210*) were then calculated for each sample and expression levels for each gene were calculated as 2^-ΔCT^ and expressed in arbitrary units.

### Histochemical assays, microscopy

For histochemical GUS assays, seedlings of five independent lines of transgenic plants were infiltrated under vacuum with GUS staining buffer: 50 mM sodium phosphate buffer (pH 7.2), 0.5 mM potassium ferrocyanide, 0.5 mM potassium ferricyanide, 10 mM EDTA, 0.01% (v/v) Triton X-100, and 1 mM 5-bromo-4-chloro-3-indolyl-glucuronic acid (X-Gluc). Incubation times ranged from 5 min to 1 h after vacuum infiltration. Seedlings were fixed in a solution of formaldehyde acetic acid ethyl alcohol (FAA) containing 10% formalin (3.7% final concentration of formaldehyde), 5% acetic acid, and 50% ethanol in water, and embedded in paraplast: 14 µm-thick tissue sections were visualised using a Nikon Eclipse Ti inverted microscope with colour CMOS camera DS-Fi3 driven by NIS software (Nikon Europe B.V., Amstelveen, The Netherlands). A 20 x/0.45 dry lens and a bright field source were used to visualise the staining. Whole seedlings of the five independent lines were imaged using scanner slides (Nanozoomer HT, Hamamatsu Photonics France, Massy, France) equipped with a 20 x lens and a bright field source.

The fluorescence patterns of WT x *proPME2::eGFP* plants, as well as those of *proCLE2::eGFP* and *proCLE4::eGFP* plants, were visualised using a Nikon Eclipse Ti inverted microscope as above. A 20 x/0.45 dry lens and a GFP B-2A filter set were used (long pass filter: excitation 450 nm/490 nm, dichroic mirror 505 nm, emission 520 nm). LecRK-I.9-tagRFP tissue localisation was visualised using a spectral confocal laser scanning system (SP8, Leica, Wetzlar, Germany) equipped with an upright microscope (DMi8, Leica). Observations were performed using an immersion lens (HC PL Fluotar N.A. 40 x/0.80, Leica). An argon laser emitting at 543 nm was used to collect fluorescence in the range between 560 and 640 nm.

### Immunohistochemistry of cell wall polysaccharides

Whole-mount immunohistochemical assays were used to analyse the occurrence of CW polysaccharides in roots using sample preparation adapted from Hejátko *et al*. (2006) and immunolabelling adapted from Francoz *et al*. (2019). The distribution patterns of polysaccharides were revealed using rat monoclonal antibodies LM19 and LM20 recognising low-methylesterified HGs and partially-methylesterified HGs, respectively (https://www.kerafast.com). Col-0 and *lecrk-I.9-1* seedlings were grown for 7 days in 250 mL Erlenmeyer flasks containing 50 mL of a half Murashige and Skoog (MS) medium (20 mM KNO_3_) (Sigma-Aldrich M5519, St Louis, MO, USA) supplemented with 1% sucrose (w/v) at pH 5.7 (KOH). Seedlings were fixed in 4 % (w/v) paraformaldehyde, 15% (v/v) dimethyl sulfoxide, 0.1 % (v/v) tween-20 in phosphate-buffered saline (PBS) solution pH 7.4 complemented with 1:1 (v/v) heptane. After three 1-min vacuum cycles, the fixation occurred for 45 min at room temperature while shaking (100 rpm). Seedlings were washed twice 5 min in 100% methanol, thrice 5 min in 100% ethanol, and then permeabilised in a solution containing ethanol and Roti®Histol (Carl Roth GmbH, Karlsruhe, Germany) (1:1, v/v, 35 min at room temperature). Seedlings were further rinsed twice in 100% ethanol and rehydrated in an ethanol series: 75%, 50% and 25% (v/v) in PBS. After four rinses in a buffer containing 10 mM Tris-HCl pH 7.5, 500 mM NaCl, 0.3% (w/v) Triton X-100 (TTBS), the seedlings were transferred to TTBS containing 5% (w/v) non-fat dry milk. Immunolabelling experiments were performed with LM19 and LM20 (1:10 dilution in TTBS-milk, 2 h incubation) followed by anti-rat IgG-Alexa488 (Invitrogen®) or anti-rat IgG alkaline phosphatase conjugate (Sigma-Aldrich A8438) secondary antibodies (1:200 dilution in TTBS-milk, 1 h incubation). Simple labelling with secondary antibodies was performed as a negative control. Alkaline phosphatase was revealed in the NBT/BCIP chromogenic substrate [nitro blue tetrazolium chloride (NBT) 350 µg.mL^-1^, 5-bromo 4-chloro-3-indolyl phosphate (BCIP) 175 µg.mL^-1^, in 100 mM Tris–HCl pH9.5, 100 mM NaCl, 10 mM MgCl_2_]. Seedlings were mounted between a slide and coverslip and imaged using a scanner (Nanozoomer HT) equipped with a 20x lens and a bright field source to visualise the blue-purple product of the enzymatic reaction with alkaline phosphatase, or with a 40x lens and a filter set (excitation 482 nm/518 nm, dichroic mirror 488 nm, emission 525 nm/530 nm) to visualise the Alexa488 fluorescence. Images were analysed using the ImageJ software (https://imagej.nih.gov/ij) to quantify the number of pixels per unit area covering the root elongation zone.

### Enzyme activity assays

*A. thaliana* seedlings from various genetic backgrounds were grown for 7 days in 250 mL Erlenmeyer flasks containing 50 mL of a half Murashige and Skoog (MS) medium (20 mM KNO_3_) (Sigma-Aldrich M5519, St Louis, MO, USA) supplemented with 1% sucrose (w/v) at pH 5.7 (KOH). Three biological replicates were conducted with one flask yielding a single biological replicate. Upon harvest, the seedlings were washed with UHQ water. For each flask, ∼1000 root segments were sampled, as for the transcriptomics study. For PME activity, a soluble protein fraction was extracted from an exact amount of tissue (a dry mass of at least 10 mg is required) by homogenisation in 400 µL of 20 mM sodium phosphate buffer pH 7.4 containing 1 M NaCl, 10 µM β-ME, 4 µL of protease inhibitor cocktail (Sigma-Aldrich P9599) and subsequently centrifuged at 12,000 *g* for 5 min at 4°C. The extraction step was repeated once and the supernatants were combined. The extracts were then desalted on Bio-Gel® P-6DG gel (BIO-RAD, Hercules, CA, USA) equilibrated with 20 mM sodium phosphate buffer pH 7.4 containing 0.15 M NaCl. PME activity was determined using the alcohol oxidase coupled assay (Klavons and Bennett, 1986). For PG activity, a similar soluble protein fraction was prepared but with 20 mM sodium acetate buffer pH 5.0. PG activity was determined by measuring the release of galacturonic acid (GalUA) residues from polygalacturonic acid (Sigma-Aldrich P3889) using GalUA as the standard (Boudart *et al*., 2003).

### Cell wall monosaccharide composition

Lyophilised root tissues were obtained as described above. Total cell wall sugar composition was determined as previously described (Lahaye *et al*., 2020). Briefly, Alcohol Insoluble Materials (AIMs) were extracted from 20 to 30 mg of lyophilised root tissues using Dionex™ ASE™ 350 accelerated solvent extractor (Thermo Fisher Scientific Inc., Waltham, MA, USA). The identification and quantification of cell wall neutral sugars were performed by gas-chromatography (Trace GC Ultra, Thermo Fisher Scientific Inc.) after sulphuric acid degradation with or without a pre-hydrolysis step using 13 M sulphuric acid to determine the amount of glucose residues from acidic-resistant cellulose. Uronic acids in acid hydrolysates were quantified using the metahydroxydiphenyl colorimetric acid method (Blumenkrantz and Asboe-Hansen, 1973).

### Degree of HG methylesterification

The degree of methylesterification was determined as follows. A dry mass of at least 10 mg was immersed in 96% boiling ethanol for 20 min. After centrifugation at 12,000 *g* for 5 min, the insoluble material was left under magnetic stirring for 15 min in 70% ethanol. This step was repeated thrice. The pellet was resuspended in 1 mL 1:1 CHCl_3_/MeOH (v/v) and left under stirring for 15 min. The residue was dried by solvent exchange (ethanol, acetone) and left overnight at room temperature to give an alcohol-insoluble residue (AIR) (Levigne *et al*., 2002). Pectins were isolated from AIR by acidic extraction (200 mM HCl pH 1.0, 95°C for 30 min). After centrifugation at 12,000 *g* for 5 min, the acidic extraction was repeated thrice. Supernatants were pooled and neutralised with 2 +N KOH. Pectins were hydrolysed in 2 N KOH at least for 30 min at room temperature and then neutralised with 2 N HCL. The uronic acid content and the degree of methylesterification were determined as described (Blumenkrantz and Asboe-Hansen, 1973; Klavons and Bennett, 1986).

### Quantification and statistical analysis

Transcriptomics and RT-qPCR data, measurements of PME and PG activities, measurements of the degree of pectin methylesterification, and counting of lateral roots per seedling were statistically analysed by Student’s t-tests. To quantify fluorescence (as defined in Fig. 3B, C), images were analysed using the ImageJ software (https://imagej.nih.gov/ij) to determine the number of pixels per unit area covering the root elongation zone. Values were statistically analysed by a Student’s t-test.

## Results

### *LecRK-I.9* is expressed at a high level in root tissues

Public transcriptomics datasets indicate a high level of expression of *LecRK-I.9* in roots (Brady *et al*., 2007; Winter *et al*., 2007). The *LecRK-I.9* tissue-specific expression pattern was determined using its promoter region fused to the β-glucuronidase (GUS) reporter (*proLecRK-I.9::GUS*). Seedlings of five independent stably transformed wild-type (WT) *A. thaliana* plants, grown in liquid culture for 7d were analysed and representative images are shown in Fig. 1. The *proLecRK-I.9* activity was mainly observed in the root apex and young tissues and decreased in the older tissues (Fig. 1A). It was also found in the collar and the phloem companion cells of the cotyledons (Fig. 1A, 1F and 1G). In the root tip, the *LecRK-I.9* promoter was active in the epidermal cells of the meristematic and elongation zones as well as in the endodermis, the pericycle and the stele, and to a lesser extent in the cortex of the meristematic zone (Fig. 1B and 1D). In differentiated tissues, it was mainly active in the endodermis, the pericycle and the stele (Fig. 1B, 1C and 1E).

**Fig. 1:**
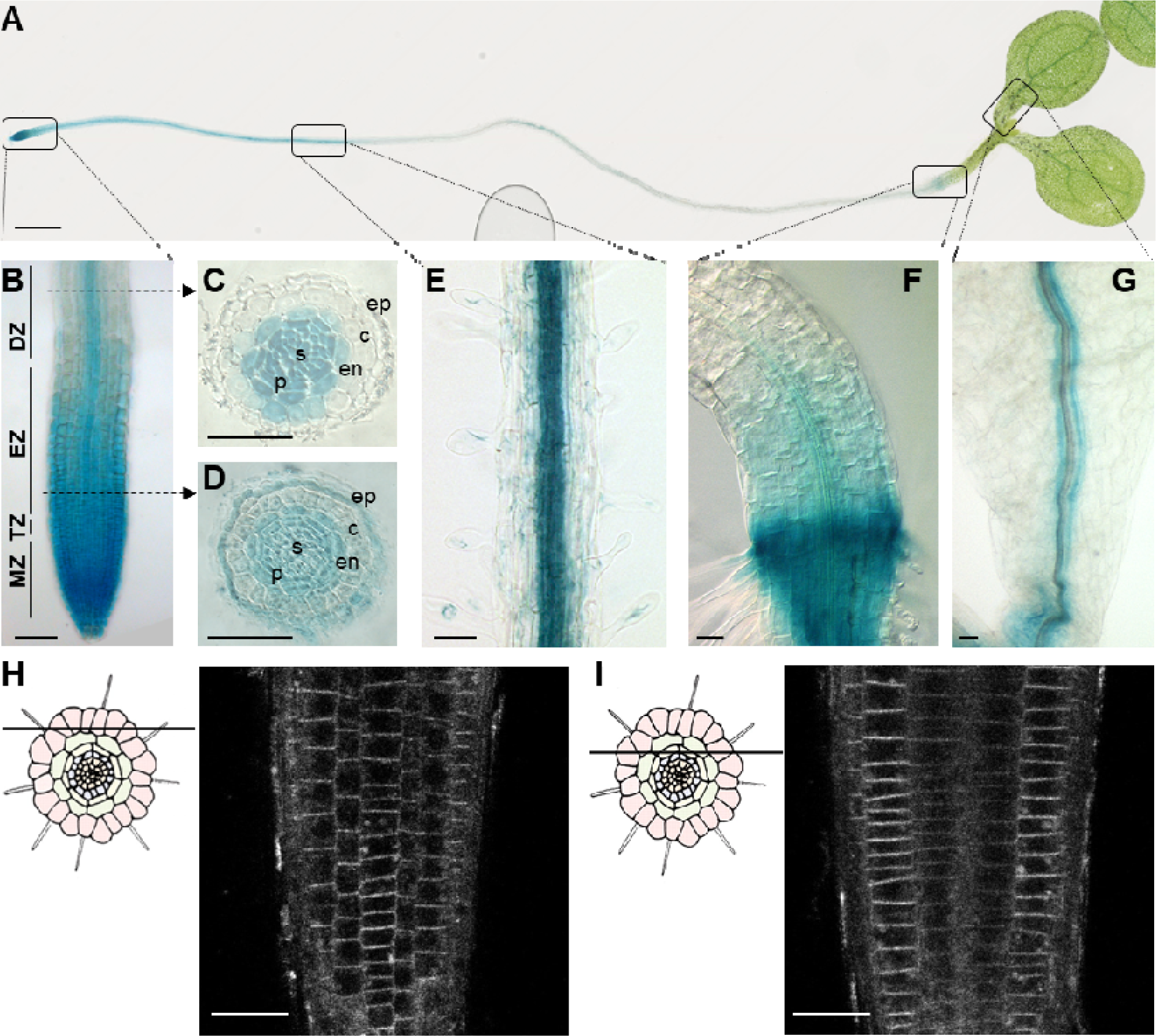
*LecRK-I.9* is mainly expressed in root tissues. (A) *pLecRK-I.9::GUS* reporter expression was detected in 7d-old seedling root tissues as well as in the collar and the veins of cotyledons. (B to G) Promoter activity of *LecRK-I.9* in a root tip (B), in root cross-sections of the differentiation zone (C) and of the beginning of the elongation zone (D), in mature root tissue (E), in collar (F) and cotyledons (G). The GUS enzymatic reaction time was 10 min. MZ = meristematic zone; EZ = elongation zone; TZ = transition zone; DZ = differentiation zone; ep = epidermis; c = cortex; en = endoderm; p = pericycle; s = stele. (H and I) Fluorescence signal in the root epidermis visualised by confocal microscopy in 7d-old root apices of *lecrk-I.9-1* complemented *proLecRK-I.9::LecRK-I.9::tagRFP A. thaliana*: optical sections through the epidermis (H) and deeper tissue layers (I) in the elongation zone. Scale bars: 2 mm (A), 50 µm (B to G), and 25 µm (H and I).

To further characterise the *LecRK-I.9* expression pattern, transgenic plants expressing the *proLecRK-I.9::LecRK-I.9::tagRFP* construct in the *lecrk-I.9-1* and *lecrk-I.9-2 loss-of-function* mutant backgrounds were generated. If the GUS activity was readily observed after a short incubation time in the previous experiment, the tagRFP fluorescence was poorly detected. It outlined the epidermis cells of the root apices (Fig. 1H and 1I) consistently with the plasma membrane localisation of LecRK-I.9 (Bouwmeester *et al*., 2011). This localisation showed a striking polarisation of the signal in the radial faces of the epidermal cells of the elongation zone. No significant signal was detected in other tissues of the seedlings. This result suggested a high turnover or a low level of expression for LecRK-I.9::tagRFP.

### Transcriptomics identifies LecRK-I.9 as a putative regulator of cell wall dynamics

*LecRK-I.9* expression was mostly localised in root tissues. The transcriptome of WT roots and *lecrk-I.9-1* roots were compared to get some insight into the functions of LecRK-I.9. Considering that the two *lecrk-I.9* mutants showed similar published phenotypes, and previous transcriptional studies were published using *lecrk-I.9-1*, the latter was chosen for the root-specific transcriptomic approach (Bouwmeester *et al*., 2011; Choi *et al*., 2014; Balagué *et al*., 2017). The differential analysis retained 20,713 genes after data processing. Three criteria of selection were used: (*i*) a mean raw value >100, (*ii*) a statistical analysis by a Student’s t-test with a *p-value <0.05*, and (*iii*) a fold-change >2.0 or <0.5 (Supplementary Dataset 1).

Thus, 199 up-regulated and 263 down-regulated genes in *lecrk-I.9-1* roots compared to WT were identified (Fig. 2; Supplementary Dataset 1). Control experiments using RT-qPCR for 17 genes confirmed the transcriptomics data (Supplementary Fig. S1). A distinctive feature of this analysis was the over-representation of genes encoding CW and CW-related proteins: 39% of the up-regulated genes (78 among 199) and 25% of the down-regulated genes (65 among 263) (Fig. 2; Supplementary Dataset 1). Most of the up-regulated genes encoded CW remodelling proteins, such as (*i*) pectin methylesterases (PMEs) and their inhibitors (PMEIs), a pectin acetylesterase (PAE) and polygalacturonases (PGs, glycoside hydrolases 28, GH28) acting on HGs; (*ii*) an expansin, a xyloglucan endotransglycosylase/hydrolase (XTH) acting on cellulose and hemicelluloses; and (iii) cellulose synthase-like proteins (CSLBs) contributing to the biosynthesis of hemicelluloses (Supplementary Dataset 1). Most of the down-regulated genes encoded proteases or proteins related to lipid metabolism such as non-specific lipid transfer proteins (nsLTPs) and GDSL esterase/lipase proteins possibly involved in cuticle formation (Supplementary Dataset 1). Besides, the genes encoding oxido-reductases and peptides were either up- or down-regulated (Fig. 2) and included CIII Prxs and plant defensin-like proteins (PDFs) encoding genes.

**Fig. 2.**
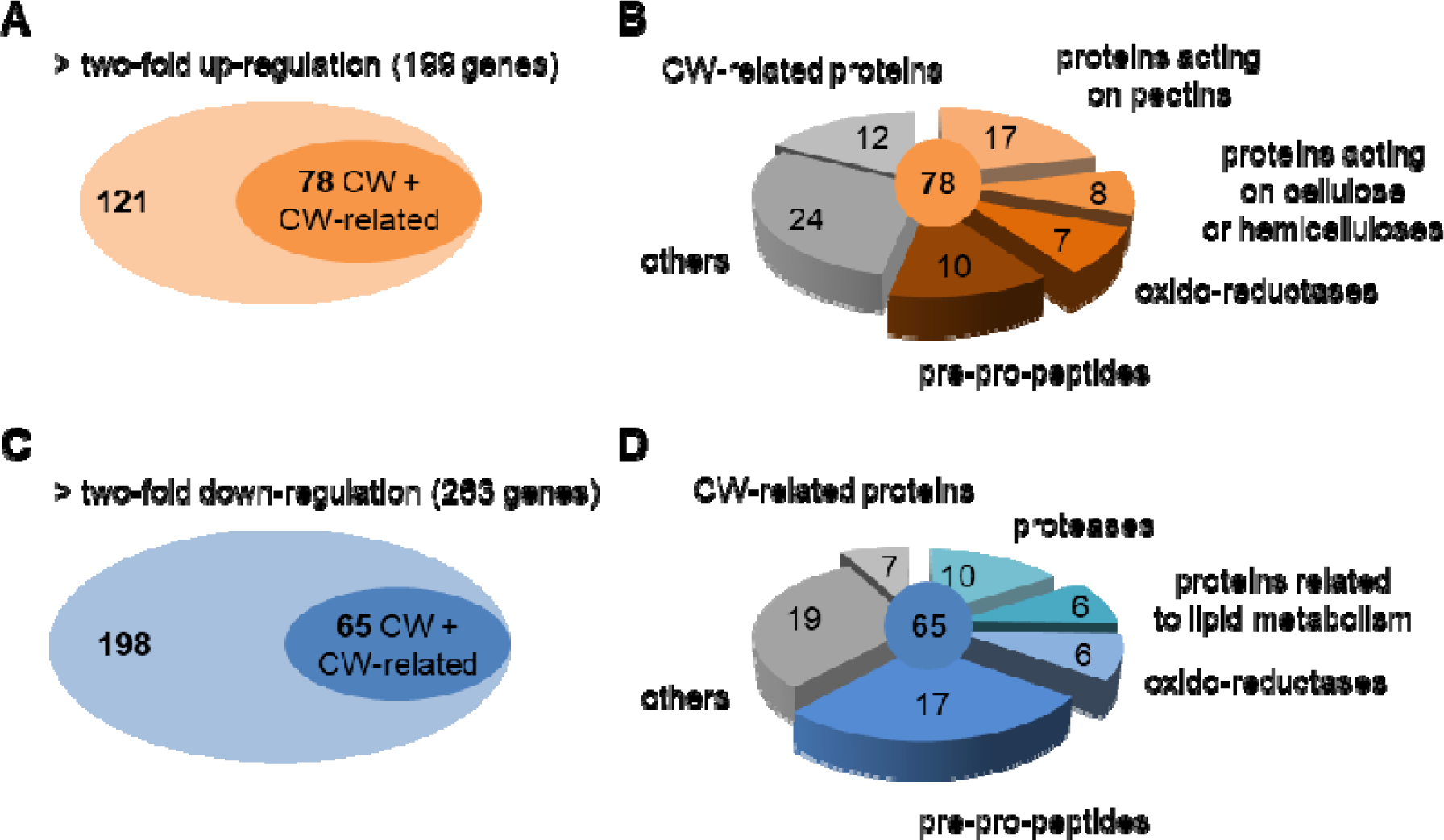
Comparative root transcriptome analysis between WT *A. thaliana* and the *lecrk-I.9-1* mutant identified genotype differentially expressed genes. Whole roots harvested from 7d-old seedlings grown in liquid culture were used for RNA isolation and microarray analysis. Genes were manually annotated to obtain a functional classification. (A) The number of genes up-regulated in *lecrk-I.9-1* with a focus on those encoding CW proteins (presence of a predicted signal peptide, absence of an intracellular retention signal) and CW-related proteins (absence of a predicted signal peptide and experimentally demonstrated CW-related function), *i.e.* 78 genes out of 199. (B) Functional classification of CW and CW-related up-regulated genes. (C) The number of genes down-regulated in *lecrk-I.9-1* with a focus on those encoding CW and CW-related proteins, *i.e.* 65 genes out of 263. (D) Functional classification of CW and CW-related down-regulated genes. The identified genes are listed in Supplementary dataset 1 with their expression ratio and locus identifier, the description of the gene, the primary gene symbol, the bibliographic references and the Student’s t-test values.

The CW-related genes encoded CW polysaccharides biosynthesis enzymes *e.g.* callose synthase complex (UDP-DEPENDENT GLYCOSYL TRANSFERASE 75B1, UGT75B1) and hemicellulose biosynthetic enzymes (XYLOGLUCAN GALACTURONOSYL-TRANSFERASE 1, XUT1) involved in root development as well as proteins involved in the development of vascular elements such as LACCASE 11 (LAC11). In addition, many genes were related to root developmental processes such as *BORON TRANSPORTER 2* (*BOR2*), *CLE2* and *CLE4*, *INDOLE-3-ACETIC ACID INDUCIBLE 27* (*IAA27*), *MIZU-KUSSEI 1* (*MIZ1*), *PME2*, *XTH21* (Supplementary Dataset 1; Supplementary Reference List). Altogether, these results suggest that LecRK-I.9 plays a role in the control of CW remodelling in roots.

The Gene Ontology term analysis (https://geneontology.org/) of the genes which are up- or down-regulated in *lecrk-I.9* confirmed the over-representation of genes encoding CW and CW-related proteins (Supplementary Table S2). The up-regulated genes revealed biological processes such as (*i*) dolichol biosynthesis that could be linked to the *N*-glycosylation pathway, (*ii*) pectic catabolism, (*iii*) CW modification and (*iv*) root development. The down-regulated genes were mostly related to biotic and abiotic constraints. In particular, the responses to bacteria and/or fungi were illustrated in previous studies (Bouwmeester *et al*., 2011; Balagué *et al*., 2017).

### *lecrk-I.9* loss-of-function mutants are altered in cell wall polysaccharides and enzyme activities

To determine whether the transcriptomics profiles correlated with changes in the root CW polysaccharide composition, the sugar contents and the degree of HG methylesterification of roots were analysed. Whole roots were collected from various genetic backgrounds: (*i*) *lecrk-I.9-1* and *lecrk-I.9-2* mutants, (*ii*) dominant negative mutants in *lecrk-I.9-1* and *lecrk-I.9-2* backgrounds (*dnm1* and *dnm2*), and (*iii*) *LecRK-I.9* over-expressor lines (*Ox-1* and *Ox-2*). The results showed no difference neither in total neutral monosaccharides, rhamnose, fucose, arabinose, xylose, mannose, or galactose, nor in uronic acid contents (Supplementary Fig. S2). Whole mount immunolocalisation was then performed using antibodies against low-methylesterified (LM19) and partially-methylesterified HGs (LM20) (Fig. 3A, B). Interestingly, a significant difference in the fluorescence intensity between *lecrk-I.9-1* and WT root apices was observed in the epidermis using LM20 antibody (mostly in EZ and DZ) (Fig. 3C), suggesting the presence of more methylesterified HGs in the *lecrk-I.9-1* epidermal cell walls. Consistently, using a biochemical assay, a higher degree of HG methylesterification was found in *lecrk-I.9-1* and *lecrk-I.9-2* as well as in *dnm-1* (Fig. 3D). The over-expressor lines *Ox-1* and *Ox-2* showed a degree of HG methylesterification similar to the WT.

**Fig. 3:**
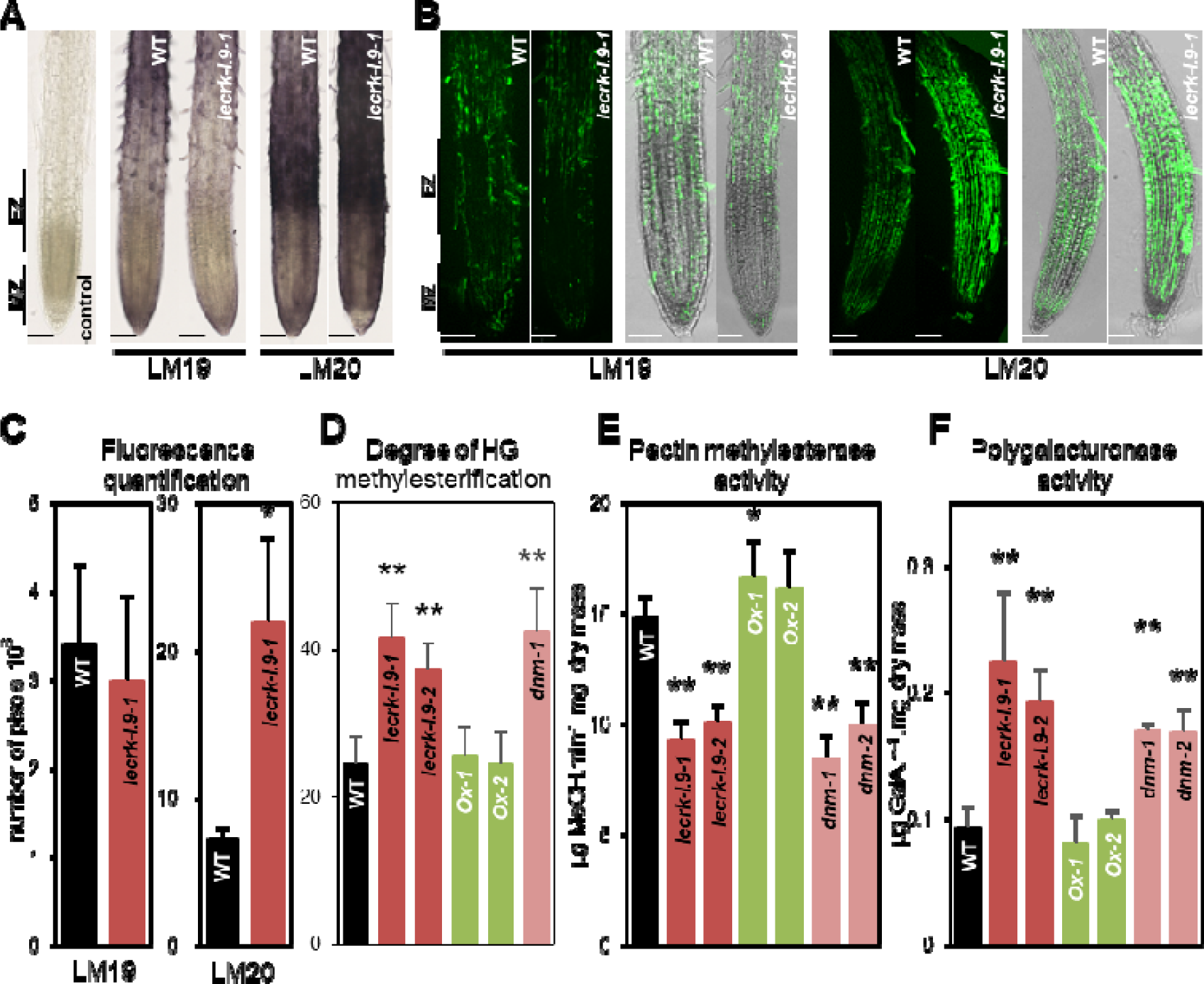
LecRK-I.9 regulates HG methylesterification and cell wall HG-related enzyme activities in root tissues. (A) and (B) Immunolocalisation of de-esterified HGs and partially methyl-esterified HGs in 7d-old seedling root tissues using LM19 and LM20 monoclonal antibodies, respectively. The meristematic zone (MZ) and elongation zone (EZ) are shown. Detection was carried out by alkaline phosphatase- (A) or alexa488 fluorochrome- (B) conjugated secondary antibodies. (B) Epifluorescence images (long pass filter, left panels) were superimposed to bright field images (right panels) for both wild-type (WT) and *lecrk-I.9-1* seedlings. Scale bars: 50 µm. (C) Quantification of the fluorescence signal obtained in (B) (n=6). (D), (E) and (F) respectively shows the degree of HG methylesterification, pectin methylesterase and polygalacturonase activities in the whole root tissues of 7d-old seedlings from different *A. thaliana* genotypes with modified *LecRK-I.9* expression levels. Ox: over-expressor lines. *dnm*: dominant negative mutant lines. The mean ± standard deviation of four biological replicates is shown. Asterisk denotes statistically significant differences according to Student’s t-test (\**P* < 0.05; *\*\*P* < 0.01) between mutant and over-expressing lines *vs* WT plants.

To probe the observed changes in CW composition, the PME and PG activities in roots were quantified. A 1.6-fold decrease in PME activity in *lecrk-I.9-1*, *lecrk-I.9-2*, *dnm-1* and *dnm-2* were detected (Fig. 3E). These results were consistent with the LM20 immunolabelling and the biochemical assay showing a higher degree of HG methylesterification in these lines. However, 7 *PMEs* were up-regulated in *lecrk-I.9-1* (Supplementary Dataset 1). Six of the 7 *PME* genes are mainly expressed in the root epidermis (*PME2*, *8*, *33*, *47*, *59* and *60*) (https://bar.utoronto.ca/eplant/), but, only one could be detected in the root proteome (*PME2*) (http://www.polebio.lrsv.ups-tlse.fr/WallProtDB/). Furthermore, two *PMEIs* (*PMEI4* and *UNE11*) expressed in the epidermis but not detected in the root proteome, were also up-regulated in *lecrk-I.9-1.* Thus, the large size of the *PME*/*PMEI* family in *A. thaliana* (66 members) and the tissue-specific gene expression could explain that the 9 up-regulated PME/PMEI genes weakly contributed to the total PME activity. These data illustrate the complex fine-tuning of the HG methylesterification status in *lecrk-I-9-1* roots. Finally, a 2-fold increase in PG activity in *lecrk-I.9-1*, *lecrk-I.9-2*, *dnm-1* and *dnm-2* compared to the WT and the over-expressor lines were detected (Fig. 3F). This was consistent with the up-regulation of several *PGs* genes in *lecrk-I.9-1* (Supplementary Dataset 1). Taken together, our data show that LecRK-I.9 plays a crucial role in remodelling HGs through the regulation of PME and PG activities.

### LecRK-I.9 negatively regulates lateral and adventitious root development

*LecRK-I.9* promoter activity was shown in the endodermis, the pericycle and the stele (Fig. 1B, 1C and 1E). CW remodelling has to occur to allow the emergence of LRs from the pericycle layer through the overlaying tissues (Banda *et al*., 2019; Swarup *et al*., 2008). Then, the number of emerged LRs and the LR initiation index were analysed (Dubrovsky & Forde, 2012). When grown in the presence of 20 mM KNO_3_ (the usual concentration of half-strength Murashige and Skoog medium), both the number of emerged LR and the LR initiation index slightly increased in *lecrk-I.9* and dominant negative mutants compared to WT, over-expressors and complemented plants (Fig. 4A, B). A lower nitrate concentration in the culture medium (KNO_3_ 0.1 mM) was also tested. Even if nitrogen has a small effect on LR number and density (Forde, 2014), it is a major element affecting CW composition and dynamics (Fernandes *et al*., 2016a; Qin *et al*., 2019; Rivai *et al*., 2021). When grown at 0.1 mM KNO_3_, *lecrk-I.9* and dominant negative mutants exhibited twice more LRs than WT, over-expressors and complemented plants (Fig. 4C). LR initiation index increased on average by 1.3 times for the *lecrk-I.9* mutants (Fig. 4D). So, both the LR initiation and the LR emergence processes were affected.

**Fig. 4:**
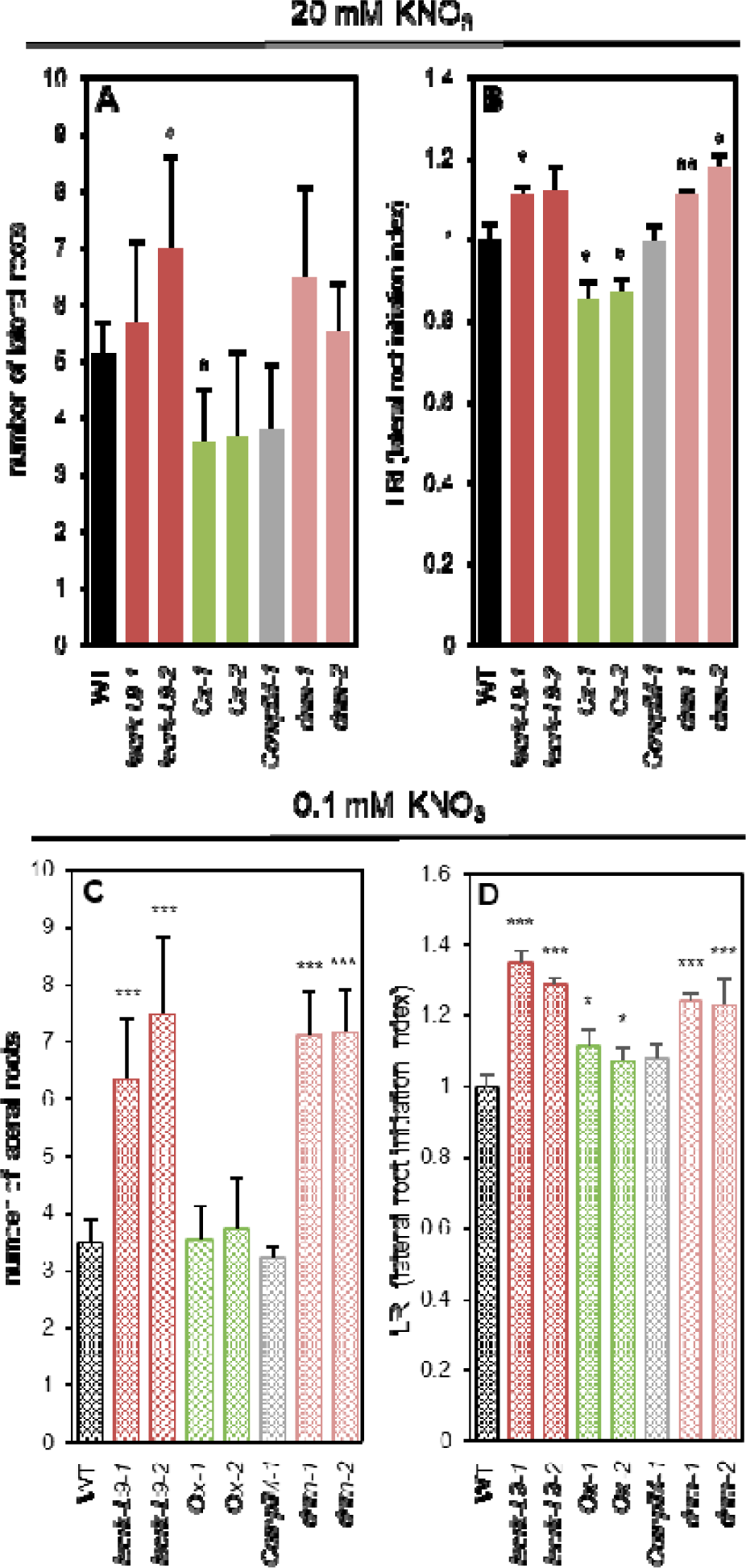
Lateral root *lecrk-I.9* phenotype. Lateral root number (A and C) and lateral root initiation index (B and D) of WT, knockout mutant (*lecrk-I.*9-1; *lecrk-I.*9-2), over-expressors (*Ox-1*; *Ox-2*), complemented (*ComplM-1*) and dominant negative mutant (*dnm-1*; *dnm-2*) lines. Seedlings were grown in two nitrate conditions (A and B, 20 mM KNO_3_; C and D, 0.1 mM KNO_3_) and observed after 11 days of culture. The mean ± standard deviation of 40<n<75 seedlings is shown. Asterisks denote statistically significant differences according to Student’s t-test (*\*\*\*P* < 0.01; *\*\*P*<0.02; *\*P*<0.05) between mutant, over-expressors, complemented and dominant negative mutant lines *vs* WT plants.

Common key regulatory elements are shared by lateral and adventitious root formation processes (Bellini *et al*., 2014). However, unlike LRs, adventitious roots emerge from non-root tissues. Here, adventitious root formation on etiolated hypocotyls was analysed. The *LecRK-I.9* promoter activity pattern observed in adventitious roots was similar to that in LRs (Supplementary Fig. S3A). As for LRs, *lecrk-I.9* mutants developed more adventitious roots than WT, over-expressors and complemented plants (Supplementary Fig. S3B).

Auxin provides a key signal during LR development (Du & Scheres, 2018). To get information about the auxin levels in the roots of the different genotypes, the auxin-responsive reporter *DR5::GUS* was introduced in these lines. Auxin accumulation was readily observed in the apex of primary roots (Supplementary Fig. S4A) and at all stages of LR development from LRPs to emerged roots (Supplementary Fig. S4B). However, no significant difference was found between the genotypes. Auxin does not appear to be targeted by the LecRK-I.9 signalling pathways.

In summary, LecRK-I.9 acts as a negative regulator of lateral and adventitious root development, independently from auxin.

### In low nitrate conditions, the LecRK-I.9 promoter activity is repressed in roots, particularly in the lateral root emergence zone

Considering the increased LR and AR phenotypes detected in low nitrate conditions, a comparative analysis of the LecRK-1.9 promoter activity and gene expression between high and low nitrate conditions was performed. Six-day-old seedlings grown in the presence of 20 mM or 0.1 mM KNO_3_ showed similar patterns of *proLecRK-I.9::GUS* expression, *i.e.* a high level of expression in the apex of the main root, and a lower one in older tissues (Fig. 5A, F). In 11d-old seedlings, fully emerged LRs were present in both culture conditions: a high promoter activity was detected in their apex as well as in their vasculature (Fig. 5B, G). However, a marked difference was observed in the seedlings when looking at LRPs. While still present at the high nitrate concentration, GUS activity was absent from LRPs at the low nitrate concentration: such a difference was observed after a short time of GUS enzymatic reaction (Fig. 5C, D, H, I). No difference was observed in fully emerged LRs (Fig. 5E, J).

**Fig. 5:**
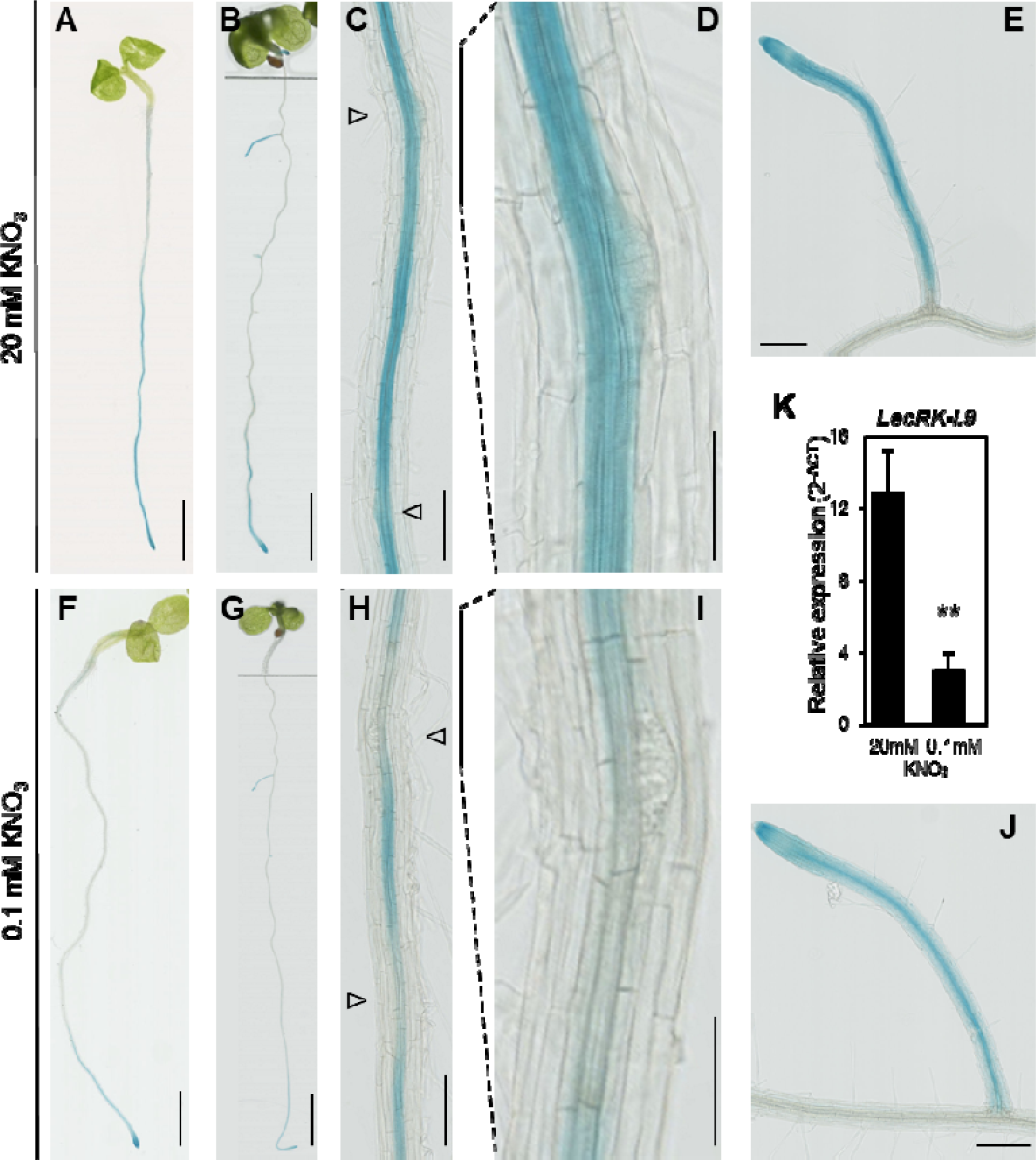
*LecRK-I.9* expression is down-regulated in roots under nitrate deficiency. *proLecRK-I.9:GUS* reporter expression was detected at both nitrate concentrations (*i.e.* 20 mM and 0,1 mM) in 6-day-old (A, F) and 11-day-old seedlings (B, G). The GUS enzymatic reaction time was 10 min. Scale bars: 5 mm (A, F) or 10 mm (B, G). (C, D, H, I) Promoter activity of *LecRK-I.9* in the zones of lateral root emergence. The GUS enzymatic reaction time was 10 min. Such a short time of reaction allows observing a lower GUS activity at the site of the lateral root primordium (Δ). This is particularly notable for the low nitrate concentration. Scale bars are 200 µm. (D, I) Close-up views of the lateral root primordia and (E, J) fully emerged lateral roots from 11d-old seedlings. Scale bars: 200 µm for emerged lateral roots; 100 µm for lateral root primordia. (K) Transcript abundance of *LecRK-I.9* was determined by RT-qPCR with cDNA generated from roots of 7-day-old wild-type seedlings grown in the two nitrate conditions. The mean ± standard deviation of five biological replicates is shown. Asterisks denote statistically significant differences according to the Student’s t-test (*\*\*P* < 0.01).

For these experiments, a lower GUS activity in the seedlings grown at the low nitrate concentration was noticed. Consistently, four times lower *LecRK-I.9* transcripts abundance in the roots of seedlings grown at 0.1 mM KNO_3_ compared to those grown at 20 mM KNO_3_ was found (Fig. 5K). Since LecRK-I.9 negatively influences the expression of genes encoding CW remodelling enzymes, the down-regulation of *LecRK-I.9* at the site of LR emergence could facilitate the emergence process.

### At both low and high nitrate concentrations, LecRK-I.9 regulates the transcription of genes encoding CLE peptide precursors involved in lateral root development

*LecRK-I.9* expression was strongly down-regulated upon nitrate deficiency. To know whether the regulation was maintained in low nitrate conditions, the level of transcripts of selected genes, such as those encoding PGs, PMEs, proteases and peptide precursors, previously shown to be up- or down-regulated in *lecrk-I.9-1* by the microarray experiment, was then measured by RT-qPCR. The expression of both up- and down-regulated selected genes in *lecrk-I.9-1* was drastically reduced at 0.1 mM KNO_3_ (Supplementary Fig. S1). For some genes up-regulated at 20 mM KNO_3_, the transcript levels were not detected (ND) at 0.1 mM KNO_3_ in both genotypes suggesting that the LecRK-I.9 control observed at 20 mM KNO_3_ was no longer effective (*e.g. GH28-1*, *PME60*). On the contrary, although the level of transcripts of *EXPA-B1* and *PME59* was much reduced in both genotypes, the regulation of their expression by LecRK-I.9 was maintained. The level of transcripts of the down-regulated genes in *lecrk-I.9-1* root transcriptome, *i.e.* those for which LecRK-I.9 acts as a positive regulator, was also reduced at 0.1 mM KNO_3_ in both genotypes, but to a lesser extent (Supplementary Fig. S1). Remarkably, the LecRK-I.9 positive regulation for *CLE2* and *CLE4* expression, was still observed under low nitrogen conditions (Fig 6A). *CLE2* and *CLE4* encode extracellular signalling peptide precursors described as negative regulators of LR development (Araya *et al*., 2014). While a strong decrease in *CLE2* and *CLE4* levels of expression in low nitrate conditions was observed, a steady state for *CLE2* and an increase in *CLE4* expression were previously reported (Araya *et al*., 2014). However, the same research group re-investigated nitrogen deficiency and showed a decrease in both *CLE2* and *CLE4* levels of expression (Ma *et al*., 2020). Also, the *CYSTEINE ENDOPEPTIDASE3 (CEP3)* gene encoding a protease homologous to a rice CEP able to digest extensins *in vitro* and induced in the context of programmed cell death (Helm *et al*., 2008), was still found to be positively regulated by LecRK-I.9.

**Fig. 6:**
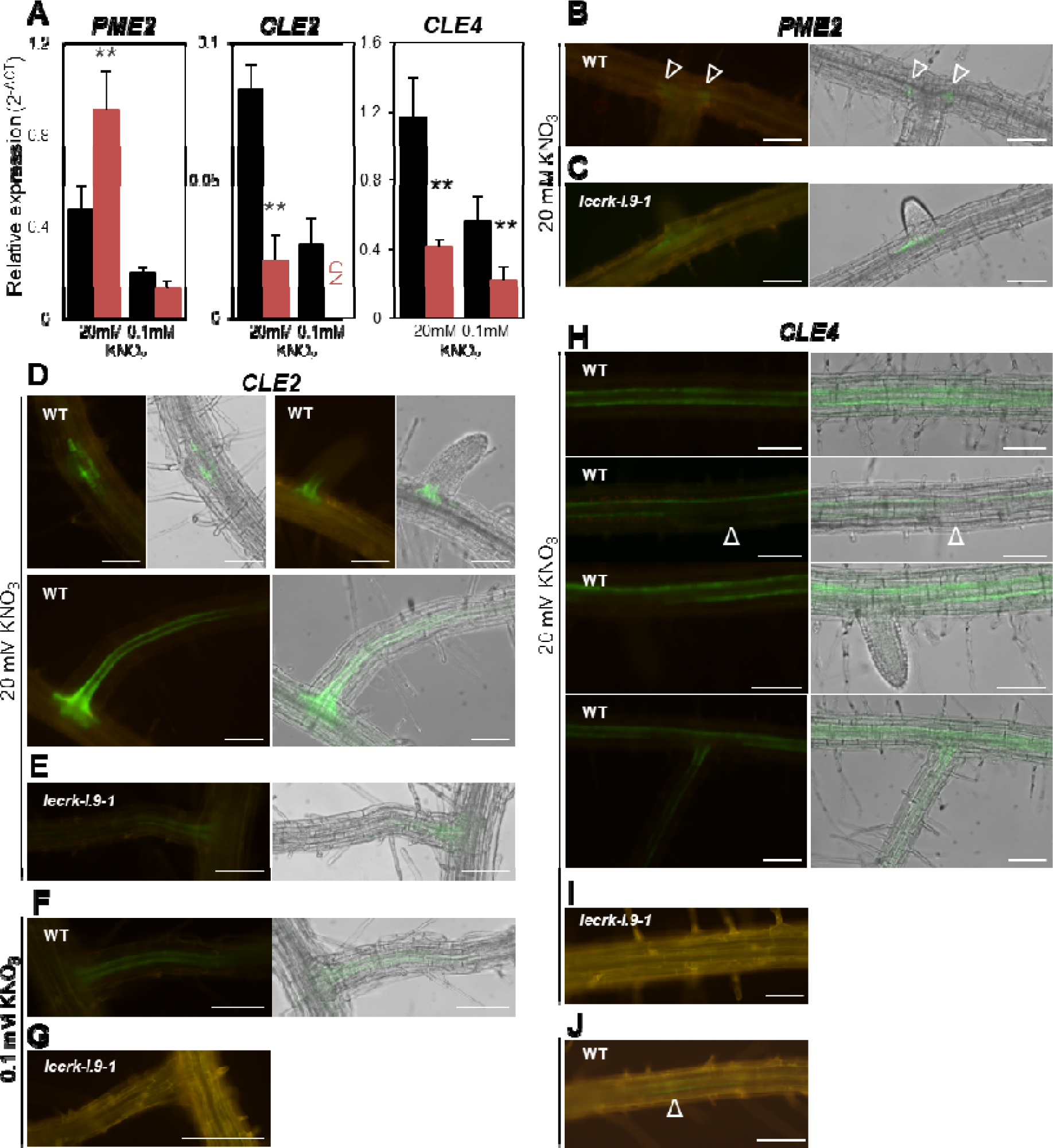
LecRK-I.9 regulates *PME2* expression at the site of lateral root emergence as well as the expression of *CLE2* and *CLE4* involved in lateral root development. (A) Transcript abundance was determined by RT-qPCR with cDNA generated from roots of 7d-old seedlings grown on a liquid medium containing either 20 mM or 0.1 mM KNO_3_: WT (black bars), *lecrk-I.9-1* (red bars). The means ± standard deviation of five biological replicates are shown. The asterisk denotes statistically significant differences according to the Student’s t-test (*\*\*P* < 0.01) between mutant lines *versus* WT plants. ND: not detected. (B, C) *proPME2:GFP* reporter expression was detected at the site of lateral root emergence in 7-day-old seedlings (20 mM KNO_3_). Epifluorescence images (long pass filter) were superimposed to bright field images for both WT (B) and mutant (*lecrk-I.9-1*) (C) seedlings. (D to G) *proCLE2:GFP* reporter expression in roots of 7d-old seedlings grown at 20 mM (D, E) or 0.1 mM KNO_3_ (F, G). (H to J) *proCLE4:GFP* reporter expression in roots of 7d-old seedlings grown at 20 mM (H, I) or 0.1 mM KNO_3_ (J). For each construct, identical microscope settings were used except for (G), (I), and (J) for which a higher exposure time was necessary due to the absence or the faintness of the fluorescent signal. Scale bars: 100 µm.

The expression patterns of *PME2*, *CLE2* and *CLE4* in roots were then examined. *PME2*, encoding a CW remodelling enzyme, was selected as a gene negatively regulated both by LecRK-I.9 and under low nitrate conditions (Fig 6A; Fig S1A). Conversely, *CLE2* and *CLE4* were selected as genes positively regulated by LecRK-I.9 and negatively regulated under low nitrate conditions (Fig 6A; Fig S1B). The promoter sequences of *PME2*, *CLE2* and *CLE4* were fused to the e*GFP* reporter and the constructs were introduced in WT and *lecrk-I.9-1* plants. A weak activity of the *PME2* promoter was revealed at 20 mM KNO_3_ in the cells adjacent to LRPs in WT (Fig. 6B) and nowhere else in the root tissues. This activity was detected early around the developing LRPs and remained identical when LRs were fully developed. In the *lecrk-I.9-1* background, the *PME2* promoter showed similar activity as in WT, but at a slightly higher level (Fig. 6C) consistently with a higher amount of *PME2* transcripts as determined by RT-qPCR (Fig. 6A).

As for *PME2*, the activity of the *CLE2* promoter at 20 mM KNO3 was detected at the early stages of LR formation in the cells surrounding LRPs (Fig. 6D), similar to the pattern previously described (Jun *et al*., 2010). As root outgrowth progressed, *CLE2* promoter activity was observed in the pericycle cells of the newly developed LRs. In fully developed LRs, the *CLE2* promoter activity was still present at the junction between the primary root and LRs but also extended in the pericycle cells without reaching the apex of LRs. In the *lecrk-I.9-1* background, a similar pattern was observed with a much lower signal intensity (Fig. 6E) underlying the strong positive regulation by LecRK-I.9. A lower intensity of the signal was also observed when the seedlings were grown at the low nitrate concentration (Fig. 6F), and no signal could be detected in the *lecrk-I.9-1* background (Fig. 6G).

The pericycle cells of the primary root of WT plants specifically displayed the eGFP reporter signal driven by the *CLE4* promoter at 20 mM KNO3 (Fig. 6H) as previously described (Jun *et al*., 2010). This signal was observed all along the primary root except in the apex (not shown) and in the zones where LRs were initiated. The eGFP signal was absent in emerged LRs and was up again in the pericycle cells of fully developed LRs. In the *lecrk-I.9-1* background, no activity of the *CLE4* promoter was detected even after long exposure (Fig. 6I). Again, it suggests that LecRK-I.9 exerts a strong positive regulation on *CLE4* expression. At the low nitrate concentration, the activity of the *CLE4* promoter was faintly observed in the pericycle cells of primary roots of WT plants (Fig. 6J), but not at all in the *lecrk-I.9-1* background (not shown).

In conclusion, LecRK-I.9 positively regulates the spatial expression of *PME2*, *CLE2* and *CLE4*. The strong positive control of the expression of these genes occurs under high nitrate conditions for *PME2* and under both high and low nitrate conditions for *CLE2* and *CLE4*.

## Discussion

The results of the present study show that LecRK-I.9, having a Legume-type lectin extracellular domain, has a developmental role in LR formation by regulating (*i*) CW remodelling enzymes at the transcriptional and post-transcriptional levels, particularly those involved in HG metabolism, and (*ii*) the transcription of *CLEs* encoding extracellular signalling peptides reported to play roles in LR formation (Araya *et al*., 2014). This is a new role since such proteins have previously been reported to be involved in the response to biotic and abiotic stresses (Bellande *et al*., 2017). In particular, LecRK-I.9 was shown to be part of the plant response to biotic and abiotic stresses (Bouwmeester *et al*., 2011; Balagué *et al*., 2017; Choi *et al*., 2014).

Our results on CW remodelling enzymes are consistent with a previous study focusing on the transcriptional response to ATP of *dorn1-1* compared to WT (Choi *et al*., 2014). *dorn1-1* carries a point mutation inactivating the kinase activity of LecRK-I.9 (Choi *et al*., 2014). Analysis of these transcriptomic data showed 23 up-regulated and 39 down-regulated genes in *dorn1-1* compared to WT (Supplementary Dataset 2). As revealed in our study, genes encoding CW proteins were prevailing, representing 35% of the up-regulated (8 among 23) and 28% of the down-regulated (11 among 39) genes. In both transcriptomic studies, the identified genes encode proteins acting on the load-bearing CW polymers (cellulose, hemicelluloses), the pectin matrix or the formation of a lignified CW (cellulose, lignin). Such a large-scale control on CW structure may explain why LecRK-I.9 was also reported to be involved in plant-pathogen interactions (Bouwmeester *et al*., 2011; Balagué *et al*., 2017), the CW being the first barrier of plant defence. The Gene Ontology term analysis confirmed the involvement of LecRK-I.9 in such processes (Supplementary Table 2).

Of particular interest is the predominance of genes encoding enzymes related to HG modifications such as PMEs and their inhibitors PMEIs, a PAE and PGs. HG, which is synthesised in the Golgi and secreted into the CW in a methylesterified and/or acetylated form, can be partially de-esterified by PMEs and PAEs in non-blockwise patterns. The resulting partially-esterified HG can be cut in shorter fragments by PGs, thus leading to CW loosening (Hocq *et al*., 2018). Alternatively, PME-dependent blockwise de-esterification of HG followed by Ca^2+^ crosslinking leads to the formation of pectin gels resulting in CW stiffening (Lamport *et al*., 2018). The pectin status influences CW structure and extensibility, which can in turn regulate organ morphogenesis (Lamport *et al*., 2018; Peaucelle *et al*., 2008; Peaucelle *et al*., 2011). Also, it was shown that a weakening or a modification of the CW of the cells overlaying LRPs, in particular the endodermis cells, is sufficient to promote their emergence (Vermeer *et al*., 2014; Roycewicz & Malamy, 2014). We show that LecRK-I.9 regulates the degree of HG methylesterification and the PME and PG enzymatic activities, consistently with the transcriptional regulation of several *PME* and *PG* genes. It also influences the balance between esterified and de-esterified HGs, as shown by our immunohistochemistry experiments. Thus, LecRK-I.9 contributes to the regulation of CW composition during secondary root organogenesis. These first outcomes have been highlighted in a model providing an overview of the regulatory network proposed for LecRK-I.9 (Fig. 7). Besides, *LecRK-I.9* expression prevails in the meristematic and elongation zones of the root apices but is also found in the pericycle and the stele of older root tissues. LecRK-I.9 may therefore function in controlling the CW dynamics at different stages of root development.

**Fig. 7:**
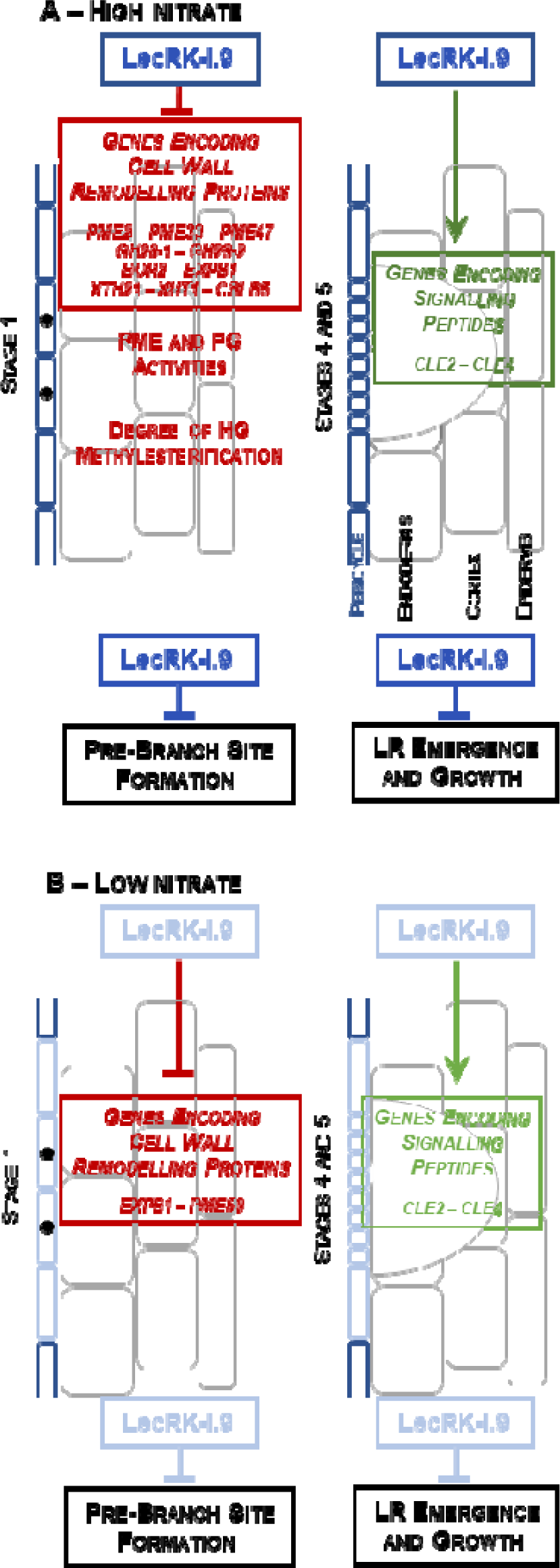
Schematic representation of the regulatory network for LecRK-I.9 in roots. LR development follows a sequence of five main stages, (1) pre-branch site formation, (2) LR initiation, (3) LR morphogenesis, (4) LR emergence, and (5) LR growth. A – Upon high nitrate conditions, *LecRK-I.9* is expressed in the pericycle (blue) and appears to be involved in CW remodelling processes at stage 1 of LR development. It is a negative regulator of genes encoding enzymes acting on pectins (*PMEs* and *PGs*), and hemicelluloses (*XTH21* and *XUT1*). The listed genes are controlled by LecRK-I.9 and are involved in the pre-branch site formation (Wachsman *et al*. 2020). Also, LecRK-I.9 regulates PME and PG activities and the degree of methylesterification of HG that is accurately controlled for the pre-branch site formation (Wachsman *et al*. 2020). On the other hand, LecRK-I.9 is a positive regulator of *CLE2* and *CLE4* encoding CW peptide precursors described as negative regulators of LR emergence and growth (stages 4 and 5) (Araya *et al*. 2014). B – Upon low nitrate conditions, the expression of *LecRK-I.9* decreases in particular at the site of LR emergence (light blue). The expression of genes encoding CW remodelling proteins is drastically reduced except *EXPB1* and *PME59*. Although the expression of positively regulated genes also decreases (light green), LecRK-I.9 maintains its regulatory function. As shown by root phenotyping, LecRK-I.9 exerts a negative regulation at stages 1, 4 and 5 of LR development.

Indeed, our phenotypic analysis shows that LecRK-I.9 is likely to have a negative role in both LR initiation and emergence processes. Recently, it was shown that both esterified and de-esterified HGs are differentially distributed at the sites of LR emergence and that genes controlling HG esterification regulate the root clock and LR initiation (Wachsman *et al*., 2020). The root clock is an oscillatory mechanism regulating gene expression, approximately every six hours, which regulates the spacing of LRs by establishing a pre-branch site (Moreno-Risueno *et al*., 2010). Wachsman *et al*. (2020) performed an RNA-seq analysis on five transverse sections including the oscillation zone, the marked pre-branch site and their flanking regions. Very interestingly, most of the genes encoding pectin-related enzymes identified in our transcriptome have their strongest expression in the oscillation zone, namely *PME2*, *PME33*, *PME47*, two PGs (*GH28-1* and *GH28-2*) and the boron transporter *BOR2*. Besides, *pme2* showed a reduced number of LRs (Wachsman *et al*., 2020). Similarly, genes encoding proteins involved in the biosynthesis or the remodelling of hemicelluloses have their strongest expression in the oscillation zone, namely those encoding the expansin *EXPB1*, the XG modifying enzymes *XTH21* and *XUT1* and the cellulose synthase-like protein *CSLB5* (Wachsman *et al*., 2020; Ramakrishna *et al*., 2019). Consistently, LecRK-I.9 appears to exert its negative regulation over genes encoding CW remodelling proteins during very early events of LR development, *i.e.* during the pre-branch site formation (Fig. 7 – stage 1) before the pre- branch site was marked.

On the other hand, LecRK-I.9 was found to be a positive regulator of the expression of genes encoding CW proteases and CW proteins related to lipid metabolism. CW proteases are likely to be involved in the turnover of CW proteins and the maturation of extracellular signalling peptides and enzymes from secreted pro-proteins. Among the CW proteins related to lipid metabolism, LTPG6 and LTP2 have been involved in the deposition of cuticular waxes (Edstam *et al*., 2013) and in the cuticle-CW interface integrity (Jacq *et al*., 2017) in aerial organs, respectively. The ABC transporter CER5 and the transcription factor MYB96 are also positively regulated by LecRK-I.9 (Supplementary Dataset 1). They are assumed to be involved in the export and biosynthesis of cuticular waxes. So, LecRK-I.9 may exert control over both the CW polysaccharides and the composition of the cuticle which has specific roles to protect and facilitate the emergence of LRs (Berhin *et al*., 2019). Additional experiments are required to precisely define the roles of the latter genes in roots.

Our study has also revealed that LecRK-I.9 has strong positive control over *CLE2* and *CLE4* expression. CLE2 and CLE4 are extracellular signalling peptides involved in LR development (Araya *et al*., 2014). Overexpression of *CLE2* and *CLE4* results in reduced LRP emergence and growth without affecting LR initiation (Araya *et al*., 2014). Our results clearly showed distinct expression patterns for *CLE2* and *CLE4*. Both *CLE4* and *LecRK-I.9* are expressed in the pericycle of lateral and primary roots and not expressed at the site of LR emergence. This is consistent with *CLE4* acting as a negative regulator of LRP emergence and growth and being positively regulated by LecRK-I.9 (Fig. 7 – stages 4 and 5). The expression pattern of *CLE2* is different, *i.e.* it is expressed in the cells surrounding LRPs and later on in the pericycle of LRs. Different functions for CLE2 and CLE4 are thus very likely.

Plant roots adapt their architecture depending on the availability of nutrients (Gruber *et al*., 2013). Among them, nitrate has received particular attention. In *A. thaliana*, nitrate exerts its effect on LR length rather than on LR number or density (Forde, 2014). However, nitrate deficiency strongly affects both the polysaccharide composition and the enzymatic repertoire of the CW (Fernandes *et al*., 2016a; Fernandes *et al*., 2016b; Qin *et al*., 2019; Rivai *et al*., 2021). All these studies showed a decrease in cellulose content which correlated with an increase in hemicellulose and/or pectin contents. Regarding the CW remodelling proteins, changes concern particular CW protein families, but also members of a given family, *e.g.* XTHs and PMEs (Qin *et al*., 2019; Rivai *et al*., 2021). From our transcriptomics experiment, the selected genes encoding CW remodelling proteins showed, at low nitrate concentration, a drastic decrease of their expression level in such a way that LecRK-I.9 only maintains its negative control on *EXPB1* and *PME59* transcription (Fig. 7B – stage 1). While *LecRK-I.9* expression was also affected (four times less), the phenotypic analysis indicated an increase in the LR initiation index in the mutant lines. It is then possible that LecRK-I.9 exerts a negative regulation on another set of CW remodelling proteins involved in the pre-branch site formation upon nitrate deficiency. In this condition, LecRK-I.9 retains its positive control on *CLE2* and *CLE4* expression to regulate LR emergence and growth (Fig. 7B – stages 4 and 5).

Auxin is a key inductive signal during LR development. Our *DR5::GUS* reporter assays showed no significant difference in auxin accumulation levels at the LR emergence sites between the tested genotypes. Nevertheless, the differential expression of genes involved in auxin transport and signal transduction were then analysed. A few of them were found: *PIN-LIKES3* (auxin transport), *MIZ1* (auxin homeostasis), *IAA27* (auxin signalling) and *GRETCHEN HAGEN3.6* (GH3.6, hormone conjugation). *MIZ1*, *IAA27* and *GH3.6* have been reported to be involved in LR development (Moriwaki *et al*., 2011; Nakazawa *et al*., 2001; Shahzad *et al*., 2020). Interestingly, *GH3.6* is also involved in the control of adventitious root formation (Gutierrez *et al*., 2012). However, the overlap between our transcriptomic data and those dedicated to auxin-dependent gene induction in LR development is extremely low (Swarup *et al*., 2008; Laskowski *et al*., 2006; Lewis *et al*., 2013). For instance, considering the genes encoding CW remodelling proteins, only *PME60* was found. Most likely, LecRK-I.9 depends on a regulation pathway that is not directly related to auxin signalling.

By regulating the expression of genes encoding several classes of proteins *e.g.* CW remodelling proteins during the pre-branch site formation and extracellular signalling peptide precursors involved in LR emergence and growth, LecRK-I.9 certainly plays a critical role in coordinating the overall LR development. However, the signalling pathway involved remains unknown. Recent studies have identified downstream targets of LecRK-I.9 in stress conditions (Chen *et al*., 2017; Kim *et al*., 2023). An important issue is to know how and which LecRK-I.9 signalling pathways are activated in developmental conditions. Since LecRK-I.9 is mediating CW-plasma membrane contacts (Gouget *et al*. 2006) and mechanical forces are causal in organ morphogenesis (Trinh *et al*., 2021), the role of LecRK-I.9 as a possible mechanosensor is a future direction of research.

## Supporting information

Supplementary reference list

## Supplementary material

- Supplementary Dataset 1. The target genes selected from the transcriptomics analysis were devoted to the comparison between WT and *lecrk-I.9-1*.
- Supplementary Dataset 2. The target genes selected from the transcriptomics analysis were devoted to the comparison between WT and *dorn1-1* (Choi *et al*. 2014).
- Supplementary Reference List. References related to the Supplementary Datasets 1 and 2.
- Supplementary Fig. S1. Control experiments using RT-qPCR to confirm the transcriptomics data.
- Supplementary Fig. S2. Monosaccharide contents of polysaccharides extracted from root tissues of different *A. thaliana* genotypes.
- Supplementary Fig. S3. Adventitious root *lecrk-I.9* phenotype.
- Supplementary Fig. S4. Auxin levels in the roots of WT and *lecrk-I.9* mutants.
- Supplementary Table S1. Oligonucleotide primers were used for cloning and RT-qPCR experiments.
- Supplementary Table S2. Gene Ontology term analysis.

## Acknowledgments

We are thankful to the Centre National de la Recherche Scientifique (CNRS) and to the Paul Sabatier-Toulouse 3 University for supporting our work. This research was funded by the TULIP LabEx project (ANR-10-LABX-41; ANR-11-IDEX-0002-02). Cell wall sugar composition was performed on the BIBS instrumental platform (http://www.bibs.inrae.fr). We also wish to thank Claude Lafitte for his help in pectin analysis, Luc Saulnier for his help on monosaccharides analysis and Aurélie Le Ru for her help with confocal microscopy images acquisitions.

## Author contributions

K.B. and H.C. designed experiments and analysed the results. K.B., D.R. and J.C. performed most of the experiments. S.LG. performed the analysis of CW polysaccharides. Y.M. and A.J. provided imaging at the microscope facilities. H.C. supervised the project. K.B., V.B., E.J. and H.C. wrote the manuscript. H.C. and E.J. managed the funding of the project. All authors read, commented on, and approved the content of the manuscript.

## Data availability

Transcriptomics data have been deposited at GEO (https://www.ncbi.nlm.nih.gov/geo) under the accession number GSE223898. Microscopy, RT-qPCR, biochemistry and plant phenotyping data reported in this paper will be shared upon request.

