## Supplementary reference list for "Receptor kinase LecRK-I.9 regulates cell wall remodelling and signalling during lateral root formation in *Arabidopsis*"

**Supplementary Reference List (**Literature cited in Supplementary Dataset 1)

### A - more abundant transcripts in *lecrk-I.9*


1 - Antisense expression of the *Arabidopsis thaliana* AtPGIP1 gene reduces polygalacturonase-inhibiting protein accumulation and enhances susceptibility to *Botrytis cinerea*. Ferrari S., Galletti R., Vairo D., Cervone F., De Lorenzo G. Mol. Plant Microbe Interact. 19:931-936(2006)

2 - Combined experimental and computational approaches reveal distinct pH dependence of pectin methylesterase inhibitors. Hocq L., Sénéchal F., Lefebvre V., Lehner A., Domon J.M., Mollet J.C., Dehors J., Pageau K., Marcelo P., Guérineau F., Kolšek K., Mercadante D., Pelloux J. Plant Physiol. 173:1075-1093(2017)

3 - Roles of BOR2, a boron exporter, in crosslinking of rhamnogalacturonan II and root elongation under boron limitation in Arabidopsis thaliana. Miwa K., Wakuta S., Takada S., Ide K., Takano J., Naito S., Omori H., Matsunaga T., Fujiwara T. Plant Physiol. 163:1699-1709(2013)

4 - Ectopic expression of Expansin3 or Expansinbeta1 causes enhanced hormone and salt stress sensitivity in Arabidopsis. Kwon Y.R., Lee H.J., Kim K.H., Hong S.W., Lee S.J., Lee H. Biotechnol. Lett. 30:1281-1288(2008)

5 - Integrative approaches to determining Csl function. Richmond T.A., Somerville C.R. Plant Mol. Biol. 47:131-143(2001)

6 - A xyloglucan endotransglucosylase/hydrolase involves in growth of primary root and alters the deposition of cellulose in Arabidopsis. Liu Y.B., Lu S.M., Zhang J.F., Liu S., Lu Y.T. Planta 226:1547-1560(2007)

7 - A galacturonic acid-containing xyloglucan is involved in Arabidopsis root hair tip growth. Pena M.J., Kong Y., York W.S., O'Neill M.A. Plant Cell 24:4511-4524(2012)

8 - A xylanase, AtXyn1, is predominantly expressed in vascular bundles, and four putative xylanase genes were identified in the Arabidopsis thaliana genome. Suzuki M., Kato A., Nagata N., Komeda Y. Plant Cell Physiol. 43:759-767(2002)

9 - Oxidation of monolignols by members of the berberine bridge enzyme family suggests a role in plant cell wall metabolism. Daniel B., Pavkov-Keller T., Steiner B., Dordic A., Gutmann A., Nidetzky B., Sensen C.W., van der Graaff E., Wallner S., Gruber K., Macheroux P. J. Biol. Chem. 290:18770-18781(2015)

10 - Laccase is necessary and nonredundant with peroxidase for lignin polymerization during vascular development in Arabidopsis. Zhao Q., Nakashima J., Chen F., Yin Y., Fu C., Yun J., Shao H., Wang X., Wang Z.Y., Dixon R.A. Plant Cell 25:3976-87(2013)

11 - IRON MAN is a ubiquitous family of peptides that control iron transport in plants. Grillet L., Lan P., Li W., Mokkapati G., Schmidt W. Nat. Plants 4:953-963(2018)

12 - The putative peptide gene FEP1 regulates iron deficiency response in Arabidopsis. Hirayama T., Lei G.J., Yamaji N., Nakagawa N., Ma J.F. Plant Cell Physiol. 59:1739-1752(2018)

13 - The Arabidopsis GASA10 gene encodes a cell wall protein strongly expressed in developing anthers and seeds. Trapalis M., Li S.F., Parish R.W. Plant Sci. 260:71-79(2017)

14 - The plant defensin gene AtPDF2.1 mediates ammonium metabolism by regulating glutamine synthetase activity in *Arabidopsis thaliana*. Yao J., Luo J.S., Xiao Y., Zhang Z. BMC Plant Biol. 19:557(2019)

15 - METACASPASE9 modulates autophagy to confine cell death to the target cells during Arabidopsis vascular xylem differentiation. Escamez S., André D., Zhang B., Bollhöner B., Pesquet E., Tuominen H. Biol. Open 5:122-129(2016)

16 - Serpin1 of *Arabidopsis thaliana* is a suicide inhibitor for metacaspase 9. Vercammen D., Belenghi B., van de Cotte B., Beunens T., Gavigan J.A., De Rycke R., Brackenier A., Inzé D., Harris J.L., van Breusegem F. J. Mol. Biol. 364:625-636(2006)

17 - Programmed cell death controlled by ANAC033/SOMBRERO determines root cap organ size in Arabidopsis. Fendrych M., Van Hautegem T., Van Durme M., Olvera-Carrillo Y., Huysmans M., Karimi M., Lippens S., Guérin C.J., Krebs M., Schumacher K., Nowack M.K. Curr. Biol. 24:931-40(2014)

18 – A novel UDP-glucose transferase is part of the callose synthase complex and interacts with phragmoplastin at the forming cell plate. Hong Z., Zhang Z., Olson J.M., Verma D.P. Plant Cell 13:769-779(2001)

19 - Identification and biochemical characterization of an Arabidopsis indole-3-acetic acid glucosyltransferase. Jackson R.G., Lim E.K., Li Y., Kowalczyk M., Sandberg G., Hoggett J., Ashford D.A., Bowles D.J. J. Biol. Chem. 276:4350-4356(2001)

20 - The activity of Arabidopsis glycosyltransferases toward salicylic acid, 4-hydroxybenzoic acid, and other benzoates. Lim E.K., Doucet C.J., Li Y., Elias L., Worrall D., Spencer S.P., Ross J., Bowles D.J. J. Biol. Chem. 277:586-592(2002)

21 - Plant pectin acetylesterase structure and function: new insights from bioinformatic analysis. Philippe F., Pelloux J., Rayon C. BMC Genomics 18:456(2017)

22 - A novel protein family directs Casparian strip formation in the endodermis. Roppolo D., De Rybel B., Denervaud Tendon V., Pfister A., Alassimone J., Vermeer J.E.M., Yamazaki M., Stierhof Y.D., Beeckman T., Geldner N. Nature 473:380-383(2011)

23 - Identification of quantitative trait loci controlling fibre length and lignin content in *Arabidopsis thaliana* stems. Capron A., Chang X.F., Hall H., Ellis B., Beatson R.P., Berleth T. J. Exp. Bot. 64:185-97(2013)

24 - Identifying new components participating in the secondary cell wall formation of vessel elements in zinnia and Arabidopsis. Endo S., Pesquet E., Yamaguchi M., Tashiro G., Sato M., Toyooka K., Nishikubo N., Udagawa-Motose M., Kubo M., Fukuda H., Demura T. Plant Cell 21:1155-1165(2009)

25 - Identification of novel proteins involved in plant cell-wall synthesis based on protein-protein interaction data. Zhou C., Yin Y., Dam P., Xu Y. J. Proteome Res. 9:5025-5037(2010)

26 - ABC transporters coordinately expressed during lignification of Arabidopsis stems include a set of ABCBs associated with auxin transport. Kaneda M., Schuetz M., Lin B.S., Chanis C., Hamberger B., Western T.L., Ehlting J., Samuels A.L. J. Exp. Bot. 62:2063-77(2011)

27 - Arabidopsis VASCULAR-RELATED UNKNOWN PROTEIN 1 regulates xylem development and growth by a conserved mechanism that modulates hormone signaling. Grienenberger E., Douglas C.J. Plant Physiol. 164:1991-2010(2014)

28 - SOMBRERO, BEARSKIN1, and BEARSKIN2 regulate root cap maturation in Arabidopsis. Bennett T., van den Toorn A., Sanchez-Perez G.F., Campilho A., Willemsen V., Snel B., Scheres B. Plant Cell 22:640-654(2010)

29 - The dynamics of root cap sloughing in Arabidopsis is regulated by peptide signalling. Shi C.L., von Wangenheim D., Herrmann U., Wildhagen M., Kulik I., Kopf A., Ishida T., Olsson V., Anker M.K., Albert M., Butenko M.A., Felix G., Sawa S., Claassen M., Friml J., Aalen R.B. Nat. Plants 4:596-604(2018)

30 - Identification of four Arabidopsis genes encoding hydroxycinnamate glucosyltransferases. Milkowski C., Baumert A., Strack D. FEBS Lett. 486:183-184(2000)

31 - Identification of glucosyltransferase genes involved in sinapate metabolism and lignin synthesis in Arabidopsis. Lim E.K., Li Y., Parr A., Jackson R., Ashford D.A., Bowles D.J. J. Biol. Chem. 276:4344-4349(2001)

32 - The hyper-fluorescent trichome phenotype of the brt1 mutant of Arabidopsis is the result of a defect in a sinapic acid: UDPG glucosyltransferase. Sinlapadech T., Stout J., Ruegger M.O., Deak M., Chapple C. Plant J. 49:655-668(2007)

33 - Activation tagging of the two closely linked genes LEP and VAS independently affects vascular cell number. van der Graaff E., Hooykaas P.J.J., Keller B. Plant J. 32:819-830(2002)

34 - Activation tagging of the LEAFY PETIOLE gene affects leaf petiole development in Arabidopsis thaliana. van der Graaff E., Dulk-Ras A.D., Hooykaas P.J.J., Keller B. Development 127:4971-4980(2000)

35 - Transcription switches for protoxylem and metaxylem vessel formation. Kubo M., Udagawa M., Nishikubo N., Horiguchi G., Yamaguchi M., Ito J., Mimura T., Fukuda H., Demura T. Genes Dev. 19:1855-1860(2005)

36 - A battery of transcription factors involved in the regulation of secondary cell wall biosynthesis in Arabidopsis. Zhong R., Lee C., Zhou J., McCarthy R.L., Ye Z.H. Plant Cell 20:2763-2782(2008)

37 - VASCULAR-RELATED NAC-DOMAIN7 is involved in the differentiation of all types of xylem vessels in Arabidopsis roots and shoots. Yamaguchi M., Kubo M., Fukuda H., Demura T. Plant J. 55:652-664(2008)

38 - VASCULAR-RELATED NAC-DOMAIN7 directly regulates the expression of a broad range of genes for xylem vessel formation. Yamaguchi M., Mitsuda N., Ohtani M., Ohme-Takagi M., Kato K., Demura T. Plant J. 66:579-590(2011)

39 - Ubiquitous and endoplasmic reticulum-located lysophosphatidyl acyltransferase, LPAT2, is essential for female but not male gametophyte development in Arabidopsis. Kim H.U., Li Y., Huang A.H.C. Plant Cell 17:1073-1089(2005)

40 - Emerging roles in plant defense for cis-jasmone-induced cytochrome P450 CYP81D11. Matthes M., Bruce T., Chamberlain K., Pickett J., Napier J. Plant Signal. Behav. 6:563-565(2011)

41 - Xenobiotic- and jasmonic acid-inducible signal transduction pathways have become interdependent at the Arabidopsis thaliana CYP81D11 promoter. Koster J., Thurow C., Kruse K., Meier A., Iven T., Feussner I., Gatz C. Plant Physiol. 159:391-402(2012)

42 - Identification of phosphomethylethanolamine N-methyltransferase from Arabidopsis and its role in choline and phospholipid metabolism. Begora M.D., Macleod M.J., McCarry B.E., Summers P.S., Weretilnyk E.A. J. Biol. Chem. 285:29147-29155(2010)

43 - The *xipotl* mutant of Arabidopsis reveals a critical role for phospholipid metabolism in root system development and epidermal cell integrity. Cruz-Ramirez A., Lopez-Bucio J., Ramirez-Pimentel G., Zurita-Silva A., Sanchez-Calderon L., Ramirez-Chavez E., Gonzalez-Ortega E., Herrera-Estrella L. Plant Cell 16:2020-2034(2004)

44 - AtNIGT1/HRS1 integrates nitrate and phosphate signals at the Arabidopsis root tip. Medici A., Marshall-Colon A., Ronzier E., Szponarski W., Wang R., Gojon A., Crawford N.M., Ruffel S., Coruzzi G.M., Krouk G. Nat. Commun. 6:6274(2015)

45 - Nicotianamine functions in the phloem-based transport of iron to sink organs, in pollen development and pollen tube growth in Arabidopsis. Schuler M., Rellan-Alvarez R., Fink-Straube C., Abadia J., Bauer P. Plant Cell 24:2380-2400(2012)

46 - *Arabidopsis thaliana* nicotianamine synthase 4 is required for proper response to iron deficiency and to cadmium exposure. Koen E., Besson-Bard A., Duc C., Astier J., Gravot A., Richaud P., Lamotte O., Boucherez J., Gaymard F., Wendehenne D. Plant Sci. 209:1-11(2013)

47 - Expression of an Arabidopsis phosphoglycerate mutase homologue is localized to apical meristems, regulated by hormones, and induced by sedentary plant-parasitic nematodes. Mazarei M., Lennon K.A., Puthoff D.P., Rodermel S.R., Baum T.J. Plant Mol. Biol. 53:513-530(2003)

48 - Discovery and analysis of cofactor-dependent phosphoglycerate mutase homologs as novel phosphoserine phosphatases in Hydrogenobacter thermophilus. Chiba Y., Oshima K., Arai H., Ishii M., Igarashi Y. J. Biol. Chem. 287:11934-11941(2012)

49 - The Arabidopsis AP2/ERF transcription factor RAP2.11 modulates plant response to low-potassium conditions. Kim M.J., Ruzicka D., Shin R., Schachtman D.P. Mol. Plant 5:1042-1057(2012)

50 - Perturbation of indole-3-butyric acid homeostasis by the UDP-glucosyltransferase UGT74E2 modulates Arabidopsis architecture and water stress tolerance. Tognetti V.B., Van Aken O., Morreel K., Vandenbroucke K., van de Cotte B., De Clercq I., Chiwocha S., Fenske R., Prinsen E., Boerjan W., Genty B., Stubbs K.A., Inze D., Van Breusegem F. Plant Cell 22:2660-2679(2010)

51 - Characterization of dehydrodolichyl diphosphate synthase of Arabidopsis thaliana, a key enzyme in dolichol biosynthesis. Cunillera N., Arro M., Fores O., Manzano D., Ferrer A. FEBS Lett. 477:170-174(2000)

52 - ELF4 is a phytochrome-regulated component of a negative-feedback loop involving the central oscillator components CCA1 and LHY. Kikis E.A., Khanna R., Quail P.H. Plant J. 44:300-313(2005)

53 - A mobile ELF4 delivers circadian temperature information from shoots to roots. Chen W.W., Takahashi N., Hirata Y., Ronald J., Porco S., Davis S.J., Nusino D.A., Kay, S.A., Mas P. Nat. Plants 6:416-426(2020)

54 - Arabidopsis Bax inhibitor-1 interacts with enzymes related to very-long-chain fatty acid synthesis. Nagano M., Kakuta C., Fukao Y., Fujiwara M., Uchimiya H., Kawai-Yamada M.J. Plant Res. 132:131-143(2019)

55 - Iron deficiency-mediated stress regulation of four subgroup Ib BHLH genes in Arabidopsis thaliana. Wang H.Y., Klatte M., Jakoby M., Bäumlein H., Weisshaar B., Bauer P. Planta 2007 226:897-908(2007)

56 - Requirement and functional redundancy of Ib subgroup bHLH proteins for iron deficiency responses and uptake in *Arabidopsis thaliana*. Wang N., Cui Y., Liu Y., Fan H., Du J., Huang Z., Yuan Y., Wu H., Ling H.Q. Mol. Plant 6:503-513(2013)

57 - FIT interacts with AtbHLH38 and AtbHLH39 in regulating iron uptake gene expression for iron homeostasis in Arabidopsis. Yuan Y., Wu H., Wang N., Li J., Zhao W., Du J., Wang D., Ling H.Q. Cell Res. 18:385-397(2008)

58 - CHL1 is a dual-affinity nitrate transporter of Arabidopsis involved in multiple phases of nitrate uptake. Liu K.H., Huang C.Y., Tsay Y.F. Plant Cell 11:865-874(1999)

59 - The Arabidopsis dual-affinity nitrate transporter gene AtNRT1.1 (CHL1) is regulated by auxin in both shoots and roots. Guo F.Q., Wang R., Crawford N.M. J. Exp. Bot. 53: 835-844(2002)

60 - CHL1 functions as a nitrate sensor in plants. Ho C.H., Lin S.H., Hu H.C., Tsay Y.F. Cell 138:1184-1194(2009)

61 - Nitrate-regulated auxin transport by NRT1.1 defines a mechanism for nutrient sensing in plants. Krouk G., Lacombe B., Bielach A., Perrine-Walker F., Malinska K., Mounier E., Hoyerova K., Tillard P., Leon S., Ljung K., Zazimalova E., Benkova E., Nacry P., Gojon A. Dev. Cell 18:927-937(2010)

62 - Multiple mechanisms of nitrate sensing by Arabidopsis nitrate transceptor NRT1.1. Bouguyon E., Brun F., Meynard D., Kubes M., Pervent M., Leran S., Lacombe B., Krouk G., Guiderdoni E., Zazimalova E., Hoyerova K., Nacry P., Gojon A. Nat. Plants 1:15015 (2015)

63 - A ferric-chelate reductase for iron uptake from soils. Robinson N.J., Procter C.M., Connolly E.L., Guerinot M.L. Nature 397:694-697(1999)

64 - Overexpression of the FRO2 ferric chelate reductase confers tolerance to growth on low iron and uncovers posttranscriptional control. Connolly E.L., Campbell N.H., Grotz N., Prichard C.L., Guerinot M.L. Plant Physiol. 133:1102-1110(2003)

65 - The FRO2 ferric reductase is required for glycine betaine's effect on chilling tolerance in Arabidopsis roots. Einset J., Winge P., Bones A.M., Connolly E.L. Physiol. Plant. 134:334-341(2008)

66 - Identification and expression analysis of a gene encoding a bacterial-type phosphoenolpyruvate carboxylase from Arabidopsis and rice. Sanchez R., Cejudo F.J. Plant Physiol. 132:949-957(2003)

67 - Structural and biochemical characterization of citrate binding to AtPPC3, a plant-type phosphoenolpyruvate carboxylase from *Arabidopsis thaliana*. Connell, M.B., Lee, M.J.Y., Li, J., Plaxton, W.C., Jia, Z. J. Struct. Biol. 204:507-512(2018)

68 - IRT1, an Arabidopsis transporter essential for iron uptake from the soil and for plant growth. Vert G., Grotz N., Dedaldechamp F., Gaymard F., Guerinot M.L., Briat J.F., Curie C. Plant Cell 14:1223-1233(2002)

69 - Regulation of iron uptake by IRT1: endocytosis pulls the trigger. Zelazny E., Vert G. Mol. Plant 8:977-979(2015)

70 - Two TIR:NB:LRR genes are required to specify resistance to Peronospora parasitica isolate Cala2 in Arabidopsis. Sinapidou E., Williams K., Nott L., Bahkt S., Toer M., Crute I., Bittner-Eddy P., Beynon J. Plant J. 38:898-909(2004)

71 - Jumonji domain protein JMJD5 functions in both the plant and human circadian systems. Jones M.A., Covington M.F., DiTacchio L., Vollmers C., Panda S., Harmer S.L. Proc. Natl. Acad. Sci. U.S.A. 107:21623-21628(2010)

72 - The essential basic helix-loop-helix protein FIT1 is required for the iron deficiency response. Colangelo E.P., Guerinot M.L. Plant Cell 16:3400-3412(2004)

73 - Salt-induced transcription factor MYB74 is regulated by the RNA-directed DNA methylation pathway in Arabidopsis. Xu R., Wang Y., Zheng H., Lu W., Wu C., Huang J., Yan K., Yang G., Zheng C. J. Exp. Bot. 66:5997-6008(2015)

74 - Phospholipase D epsilon and phosphatidic acid enhance Arabidopsis nitrogen signaling and growth. Hong Y., Devaiah S.P., Bahn S.C., Thamasandra B.N., Li M., Welti R., Wang X. Plant J. 58:376-387(2009)

75 - A conserved core of programmed cell death indicator genes discriminates developmentally and environmentally induced programmed cell death in plants. Olvera-Carrillo Y., Van Bel M., Van Hautegem T., Fendrych M., Huysmans M., Simaskova M., van Durme M., Buscaill P., Rivas S., Coll N.S., Coppens F., Maere S., Nowack M.K. Plant Physiol. 169:2684-99(2015)

76 - A novel putative auxin carrier family regulates intracellular auxin homeostasis in plants. Barbez E., Kubes M., Rolcik J., Beziat C., Pencik A., Wang B., Rosquete M.R., Zhu J., Dobrev P.I., Lee Y., Zazimalova E., Petrasek J., Geisler M., Friml J., Kleine-Vehn J. Nature 485:119-122(2012)

77 - The genetic basis of constitutive and herbivore-induced ESP-independent nitrile formation in Arabidopsis. Burow M., Losansky A., Müller R., Plock A., Kliebenstein D.J., Wittstock U. Plant Physiol. 149:561-574(2009)

78 - Blue light- and low temperature-regulated COR27 and COR28 play roles in the Arabidopsis circadian clock. Li X., Ma D., Lu S.X., Hu X., Huang R., Liang T., Xu T., Tobin E.M., Liu H. Plant Cell 28:2755-2769(2016)

79 - Arabidopsis glutathione transferases U24 and U25 exhibit a range of detoxification activities with the environmental pollutant and explosive, 2,4,6-trinitrotoluene. Gunning V., Tzafestas K., Sparrow H., Johnston E.J., Brentnall A.S., Potts J.R., Rylott E.L., Bruce N.C. Plant Physiol. 165:854-865(2014)

80 - STRESS INDUCED FACTOR 2, a leucine-rich repeat kinase regulates basal plant pathogen defense. Yuan N., Yuan S., Li Z., Zhou M., Wu P., Hu Q., Wang L., Mendu V., Luo H. Plant Physiol. 176:3062-3080(2018)

81 - Plant RNases T2, but not Dicer-like proteins, are major players of tRNA-derived fragments biogenesis. Megel C., Hummel G., Lalande S., Ubrig E., Cognat V., Morelle G., Salinas-Giegé T., Duchêne A.M., Maréchal-Drouard L. Nucleic Acids Res. 47:941-952(2019)

82 - A promoter-swap strategy between the AtALMT and AtMATE genes increased Arabidopsis aluminum resistance and improved carbon-use efficiency for aluminum resistance. Liu J., Luo X., Shaff J., Liang C., Jia X., Li Z., Magalhaes J., Kochian L.V. Plant J. 71:327-337(2012)

83 - CYSTM, a novel non-secreted cysteine-rich peptide family, involved in environmental stresses in *Arabidopsis thaliana*. Xu Y., Yu Z., Zhang D., Huang J., Wu C., Yang G., Yan K., Zhang S., Zheng C. Plant Cell Physiol. 59:423-438(2018)

84 - Malonylation is a key reaction in the metabolism of xenobiotic phenolic glucosides in Arabidopsis and tobacco. Taguchi G., Ubukata T., Nozue H., Kobayashi Y., Takahi M., Yamamoto H., Hayashida N. Plant J. 63:1031-1041(2010)

85 - STOP2 activates transcription of several genes for Al- and low pH-tolerance that are regulated by STOP1 in Arabidopsis. Kobayashi Y., Ohyama Y., Kobayashi Y., Ito H., Iuchi S., Fujita M., Zhao C.R., Tanveer T., Ganesan M., Kobayashi M., Koyama H. Mol. Plant 7:311-322(2014)

86 - LIFEGUARD proteins support plant colonization by biotrophic powdery mildew fungi. Weis C., Huckelhoven R., Eichmann R. J. Exp. Bot. 64:3855-67(2013)

87 - Myrosinases from root and leaves of *Arabidopsis thaliana* have different catalytic properties. Andersson D., Chakrabarty R., Bejai S., Zhang J., Rask L., Meijer J. Phytochemistry 70:1345-1354(2009)

88 - Glucosinolate metabolism and its control. Grubb C.D., Abel S. Trends Plant Sci. 11:89-100(2006)

89 - Arabidopsis myrosinase genes AtTGG4 and AtTGG5 are root-tip specific and contribute to auxin biosynthesis and root-growth regulation. Fu L., Wang M., Han B., Tan D., Sun X., Zhang J. Int. J. Mol. Sci. 17:E892(2016)

90 - Altered profile of secondary metabolites in the root exudates of Arabidopsis ATP-binding cassette transporter mutants. Badri D.V., Loyola-Vargas V.M., Broeckling C.D., De-la-Peña C., Jasinski M., Santelia D., Martinoia E., Sumner L.W., Banta L.M., Stermitz F., Vivanco J.M. Plant Physiol. 146:762-71(2008)

91 - MOTHER OF FT AND TFL1 regulates seed germination through a negative feedback loop modulating ABA signaling in Arabidopsis. Xi W., Liu C., Hou X., Yu H. Plant Cell 22:1733-1748(2010)

92 - MOTHER-OF-FT-AND-TFL1 represses seed germination under far-red light by modulating phytohormone responses in Arabidopsis thaliana. Vaistij F.E., Barros-Galvão T., Cole A.F., Gilday A.D., He Z., Li Y., Harvey D., Larson T.R., Graham I.A. Proc. Natl. Acad. Sci. U.S.A. 115:8442-8447(2018)

93 - ALD1 regulates basal immune components and early inducible defense responses in Arabidopsis. Cecchini N.M., Jung H.W., Engle N., Tschaplinski T.J., Greenberg J. Mol. Plant Microbe Interact. 28:455-466(2015)

94 - A gene regulatory network for root epidermis cell differentiation in Arabidopsis. Bruex A., Kainkaryam R.M., Wieckowski Y., Kang Y.H., Bernhardt C., Xia Y., Zheng X., Wang J.Y., Lee M.M., Benfey P., Woolf P.J., Schiefelbein J. PLoS Genet. 8:e1002446(2012)

95 - Cis-element- and transcriptome-based screening of root hair-specific genes and their functional characterization in Arabidopsis. Won S.K., Lee Y.J., Lee H.Y., Heo Y.K., Cho M., Cho H.T. Plant Physiol. 150:1459-1473(2009)

96 - Expression, localisation and phylogeny of a novel family of plant-specific membrane proteins. Kasaras A., Kunze R. Plant Biol. (Stuttg) 12:140-152(2010)

97 - Arabidopsis glycosyltransferases as biocatalysts in fermentation for regioselective synthesis of diverse quercetin glucosides. Lim E.K., Ashford D.A., Hou B., Jackson R.G., Bowles D.J. Biotechnol. Bioeng. 87:623-631(2004)

98 - Pathogen-responsive expression of glycosyltransferase genes UGT73B3 and UGT73B5 is necessary for resistance to Pseudomonas syringae pv tomato in Arabidopsis. Langlois-Meurinne M., Gachon C.M., Saindrenan P. Plant Physiol. 139:1890-1901(2005)

99 - Identification and evolution of functional alleles of the previously described pollen specific myrosinase pseudogene AtTGG6 in Arabidopsis thaliana. Fu L., Han B., Tan D., Wang M., Ding M., Zhang J. Int. J. Mol. Sci. 17:262(2016)

100 - Arabidopsis glucosyltransferases with activities toward both endogenous and xenobiotic substrates. Messner B., Thulke O., Schaeffner A.R. Planta 217:138-146(2003)

101 - Transcriptional and posttranscriptional regulation of transcription factor expression in Arabidopsis roots. Lee J.Y., Colinas J., Wang J.Y., Mace D., Ohler U., Benfey P.N. Proc. Natl. Acad. Sci. U.S.A. 103:6055-6060(2006)

102 - Low phosphate activates STOP1-ALMT1 to rapidly inhibit root cell elongation. Balzergue C., Dartevelle T., Godon C., Laugier E., Meisrimler C., Teulon J.M., Creff A., Bissler M., Brouchoud C., Hagege A., Muller J., Chiarenza S., Javot H., Becuwe-Linka N., David P., Peret B., Delannoy E., Thibaud M.C., Armengaud J., Abel S., Pellequer J.L., Nussaume L., Desnos T. Nat. Commun. 8:15300(2017)

103 - The FANTASTIC FOUR proteins influence shoot meristem size in *Arabidopsis thaliana*. Wahl V., Brand L.H., Guo Y.L., Schmid M. BMC Plant Biol. 10:285-285(2010)

104 - Polyprenols are synthesized by a plastidial cis-prenyltransferase and influence photosynthetic performance. Akhtar T.A., Surowiecki P., Siekierska H., Kania M., Van Gelder K., Rea K.A., Virta L.K.A., Vatta M., Gawarecka K., Wojcik J., Danikiewicz W., Buszewicz D., Swiezewska E., Surmacz L. Plant Cell 29:1709-1725(2017)

105 - A gene essential for hydrotropism in roots. Kobayashi A., Takahashi A., Kakimoto Y., Miyazawa Y., Fujii N., Higashitani A., Takahashi H. Proc. Natl. Acad. Sci. U.S.A. 104:4724-4729(2007)

106 - Hormonal regulation of lateral root development in Arabidopsis modulated by MIZ1 and requirement of GNOM activity for MIZ1 function. Moriwaki T., Miyazawa Y., Kobayashi A., Uchida M., Watanabe C., Fujii N., Takahashi H. Plant Physiol. 157:1209-1220(2011)

107 - MIZ1 regulates ECA1 to generate a slow, long-distance phloem-transmitted Ca2+ signal essential for root water tracking in Arabidopsis. Shkolnik D., Nuriel R., Bonza M.C., Costa A., Fromm H. Proc. Natl. Acad. Sci. U.S.A. 115:8031-8036(2018)

108 - SIEVE ELEMENT-LINING CHAPERONE1 restricts aphid feeding on Arabidopsis during heat stress. Kloth K.J., Busscher-Lange J., Wiegers G.L., Kruijer W., Buijs G., Meyer R.C., Albrectsen B.R., Bouwmeester H.J., Dicke M., Jongsma M.A. Plant Cell 29:2450-2464(2017)

109 - Integration of ovular signals and exocytosis of a Ca2+ channel by MLOs in pollen tube guidance. Meng J.G., Liang L., Jia P.F., Wang Y.C., Li H.J., Yang W.C. Nat. Plants 6:143-153(2020)

110 - Identification of BFN1, a bifunctional nuclease induced during leaf and stem senescence in Arabidopsis. Perez-Amador M.A., Abler M.L., De Rocher E.J., Thompson D.M., van Hoof A., LeBrasseur N.D., Lers A., Green P.J. Plant Physiol. 122:169-179(2000)

111 - Expression analysis of the BFN1 nuclease gene promoter during senescence, abscission, and programmed cell death-related processes. Farage-Barhom S., Burd S., Sonego L., Perl-Treves R., Lers A. J. Exp. Bot. 59:3247-3258(2008)

276 – Cell wall remodeling and vesicle trafficking mediate the root clock in Arabidopsis. Wachsman G., Zhang J., Moreno-Risueno M.A., Anderson C.T., Benfey P.N. Science 370:819-823(2020)

**B - less abundant transcripts in *lecrk-I.9***


112 - The MYB80 transcription factor is required for pollen development and the regulation of tapetal programmed cell death in *Arabidopsis thaliana*. Phan H.A., Iacuone S., Li S.F., Parish R.W. Plant Cell 23:2209-2224(2011)

113 - Autophagy mitigates high-temperature injury in pollen development of *Arabidopsis thaliana*. Dündar G., Shao Z., Higashitani N., Kikuta M., Izumi M., Higashitani A. Dev. Biol. 456:190-200(2019)

114 - Cysteine protease enhances plant-mediated bollworm RNA interference. Mao Y.B., Xue X.Y., Tao X.Y., Yang C.Q., Wang L.J., Chen X.Y. Plant Mol. Biol. 83:119-129(2013)

115 - KDEL-tailed cysteine endopeptidases involved in programmed cell death, intercalation of new cells, and dismantling of extensin scaffolds. Helm M., Schmid M., Hierl G., Terneus K., Tan L., Lottspeich F., Kieliszewski M.J., Gietl C. Am. J. Bot. 95:1049-1062(2008)

117 - Copper amine oxidase 8 regulates arginine-dependent nitric oxide production in *Arabidopsis thaliana*. Groß F., Rudolf E.E., Thiele B., Durner J., Astier J. J. Exp. Bot. 68:2149-2162(2017)

118 - Cross-talk between reactive oxygen species and polyamines in regulation of ion transport across the plasma membrane: implications for plant adaptive responses. Pottosin I., Velarde-Buendía A.M., Bose J., Zepeda-Jazo I., Shabala S., Dobrovinskaya O. J. Exp. Bot. 65:1271-1283(2014).

119 - Copper-containing amine oxidases contribute to terminal polyamine oxidation in peroxisomes and apoplast of *Arabidopsis thaliana*. Planas-Portell J., Gallart M., Tiburcio A.F., Altabella T. BMC Plant Biol. 13:109-109(2013)

120 - Natural variation reveals a key role for rhamnogalacturonan I in seed outer mucilage and underlying genes. Fabrissin I., Cueff G., Berger A., Granier F., Salle C., Poulain D., Ralet M.C., North H.M. Plant Physiol. 181:1498-1518(2019)

121 - Two cell wall associated peroxidases from Arabidopsis influence root elongation. Passardi F., Tognolli M., De Meyer M., Penel C., Dunand C. Planta 223:965-974(2006)

122 - The apoplastic oxidative burst peroxidase in Arabidopsis is major component of pattern-triggered immunity. Daudi A., Cheng Z., O'Brien J.A., Mammarella N., Khan S., Ausubel F.M., Bolwell G.P. Plant Cell 24:275-287(2012)

123 - Contribution of cell wall peroxidase- and NADPH oxidase -derived reactive oxygen species to *Alternaria brassicicola*-induced oxidative burst in Arabidopsis. Kámán-Tóth E., Dankó T., Gullner G., Bozsó Z., Palkovics L., Pogány M. Mol. Plant Pathol. 20:485-499(2019)

124 - The Arabidopsis lipid transfer protein 2 (AtLTP2) is involved in cuticle-cell wall interface integrity and in etiolated hypocotyl permeability. Jacq A., Pernot C., Martinez Y., Domergue F., Payré B., Jamet E., Burlat V., Pacquit V. B. Front. Plant Sci. 8:263 (2017)

125 – Coexpression patterns indicate that GPI-anchored non-specific lipid transfer proteins are involved in accumulation of cuticular wax, suberin and sporopollenin. Edstam M.M., Blomqvist K., Eklöf A., Wennergren U., Edqvist J. Plant Mol. Biol. 83:625-649(2013)

126 - Involvement of lipid transfer proteins in resistance against a non-host powdery mildew in *Arabidopsis thaliana*. Fahlberg P., Buhot N., Johansson O.N., Andersson M.X. Mol. Plant Pathol. 20:69-77(2019)

127 - CLE-CLAVATA1 peptide-receptor signaling module regulates the expansion of plant root systems in a nitrogen-dependent manner. Araya T, Miyamoto M, Wibowo J, Suzuki A, Kojima S, Tsuchiya YN, Sawa S, Fukuda H, von Wirén N, Takahashi H. Proc. Natl. Acad. Sci. U.S.A. 111:2029-2034(2014)

128 - CLE peptide signaling and crosstalk with phytohormones and environmental stimuli. Wang G., Zhang G., Wu M. Front. Plant Sci. 6:1211(2016)

129 - The secreted peptide PIP1 amplifies immunity through receptor-like kinase 7. Hou S., Wang X., Chen D., Yang X., Wang M., Turra D., Di Pietro A., Zhang W. PLoS Pathog. 10:e1004331(2014)

130 - PAMP-induced peptide 1 cooperates with salicylic acid to regulate stomatal immunity in Arabidopsis thaliana. Hou S., Shen H., Shao H. Plant Signal. Behav. 17:1666657(2019)

131 - Wound-induced polypeptides improve resistance against *Pseudomonas syringae* pv. tomato DC3000 in Arabidopsis. Yu L., Wang Y., Liu Y., Li N., Yan J., Luo L. Biochem. Biophys. Res. Commun. 504:149-156(2018)

132 - IDL6-HAE/HSL2 impacts pectin degradation and resistance to *Pseudomonas syringae* pv tomato DC3000 in Arabidopsis leaves. Wang X., Hou S., Wu Q., Lin M., Acharya B.R., Wu D., Zhang W. Plant J. 89:250-263(2017)

133 - Identification of a role for an E6-like 1 gene in early pollen-stigma interactions in *Arabidopsis thaliana*. Doucet J., Truong C., Frank-Webb E., Lee H.K., Daneva A., Gao Z., Nowack M.K., Goring, D.R. Plant Reprod. 32:307-322(2019)

134 - Cytokinin-deficient transgenic Arabidopsis plants show multiple developmental alterations indicating opposite functions of cytokinins in the regulation of shoot and root meristem activity. Werner T., Motyka V., Laucou V., Smets R., Van Onckelen H., Schmulling T. Plant Cell 15:2532-2550(2003)

135 - Cytokinin as a mediator for regulating root system architecture in response to environmental cues. Ramireddy E., Chang L., Schmülling T. Plant Signal. Behav. 9:e27771(2014)

136 - Extracellular peptide Kratos restricts cell death during vascular development and stress in Arabidopsis. Escamez S., Stael S., Vainonen J., Willems P., Jin H., Kimura S., Van Breusegem F., Gevaert K., Wrzaczek M., Tuominen H. J. Exp. Bot. 70:2199-2210(2019)

137 - Identification of a cis-regulatory element for L1 layer-specific gene expression, which is targeted by an L1-specific homeodomain protein. Abe M., Takahashi T., Komeda Y. Plant J. 26:487-494(2001)

138 - Formation of the stomatal outer cuticular ledge requires a guard cell wall proline-rich protein. Hunt L., Amsbury S., Baillie A., Movahedi M., Mitchell A., Afsharinafar M., Swarup K., Denyer T., Hobbs J.K., Swarup R., Fleming A.J., Gray J.E. Plant Physiol. 174:689-699(2017)

139 - AtBXL1, a novel higher plant (*Arabidopsis thaliana*) putative beta-xylosidase gene, is involved in secondary cell wall metabolism and plant development. Goujon T., Minic Z., El Amrani A., Lerouxel O., Aletti E., Lapierre C., Joseleau J.P., Jouanin L. Plant J. 33:677-90(2003)

140 - Functional genomic analysis of *Arabidopsis thaliana* glycoside hydrolase family 1. Xu Z., Escamilla-Trevino L.L., Zeng L., Lalgondar M., Bevan D.R., Winkel B.S.J., Mohamed A., Cheng C.L., Shih M.C., Poulton J.E., Esen A. Plant Mol. Biol. 55:343-367(2004)

141 - AtPep3 is a hormone-like peptide that plays a role in the salinity stress tolerance of plants. Nakaminami K., Okamoto M., Higuchi-Takeuchi M., Yoshizumi T., Yamaguchi Y., Fukao Y., Shimizu M., Ohashi C., Tanaka M., Matsui M., Shinozaki K., Seki M., Hanada K. Proc. Natl. Acad. Sci. U.S.A. 115:5810-5815(2018)

142 - The plant cell wall integrity maintenance and immune signaling systems cooperate to control stress responses in *Arabidopsis thaliana*. Engelsdorf T., Gigli-Bisceglia N., Veerabagu M., McKenna J.F., Vaahtera L., Augstein F., Van der Does D., Zipfel C., Hamann T. Sci. Signal. 11: 3070(2018)

143 - Evidence for a role of AtCAD 1 in lignification of elongating stems of *Arabidopsis thaliana*. Eudes A., Pollet B., Sibout R., Do C.-T., Seguin A., Lapierre C., Jouanin L. Planta 225:23-39(2006)

144 - SG2-type R2R3-MYB transcription factor MYB15 controls defense-induced lignification and basal immunity in Arabidopsis. Chezem W.R., Memon A., Li F.S., Weng J.K., Clay N.K. Plant Cell 29:1907-1926(2017)

145 - Two cinnamoyl-CoA reductase (CCR) genes from *Arabidopsis thaliana* are differentially expressed during development and in response to infection with pathogenic bacteria. Lauvergeat V., Lacomme C., Lacombe E., Lasserre E., Roby D., Grima-Pettenati J. Phytochemistry 57:1187-1195(2001)

146 - Identification of novel proteins involved in plant cell-wall synthesis based on protein-protein interaction data. Zhou C., Yin Y., Dam P., Xu Y.J. Proteome Res. 9:5025-5037(2010)

147 - CYCLIN-DEPENDENT KINASE G1 is associated with the spliceosome to regulate CALLOSE SYNTHASE5 splicing and pollen wall formation in Arabidopsis. Huang X.Y., Niu J., Sun M.X., Zhu J., Gao J.F., Yang J., Zhou Q., Yang Z.N. Plant Cell 25:637-648(2013)

148 - Characterization of the NRT2.6 gene in *Arabidopsis thaliana*: a link with plant response to biotic and abiotic stress. Dechorgnat J., Patrit O., Krapp A., Fagard M., Daniel-Vedele F. PLoS ONE 7:e42491(2012)

149 - The NRT2.5 and NRT2.6 genes are involved in growth promotion of Arabidopsis by the plant growth-promoting rhizobacterium (PGPR) strain *Phyllobacterium brassicacearum* STM196. Kechid M., Desbrosses G., Rokhsi W., Varoquaux F., Djekoun A., Touraine B. New Phytol. 198:514-524(2013)

150 - A novel role for oleosins in freezing tolerance of oilseeds in *Arabidopsis thaliana*. Shimada T.L., Shimada T., Takahashi H., Fukao Y., Hara-Nishimura I. Plant J. 55:798-809(2008)

151 - Specialization of oleosins in OB dynamics during seed development in Arabidopsis thaliana seeds. Miquel M., Trigui G., D'Andréa S., Kelemen Z., Baud S., Berger A., Deruyffelaere C., Trubuil A., Lepiniec L., Dubreucq B. Plant Physiol. 164:1866-78(2014)

152 - Bet v 1 from birch pollen is a lipocalin-like protein acting as allergen only when devoid of iron by promoting Th2 lymphocytes. Roth-Walter F., Gomez-Casado C., Pacios L.F., Mothes-Luksch N., Roth G.A., Singer J., Diaz-Perales A., Jensen-Jarolim E. J. Biol. Chem. 289:17416-17421(2014)

153 - The HECATE genes regulate female reproductive tract development in *Arabidopsis thaliana*. Gremski K., Ditta G., Yanofsky M.F. Development 134:3593-3601(2007)

154 - Arabidopsis HECATE genes function in phytohormone control during gynoecium development. Schuster C., Gaillochet C., Lohmann J.U. Development 142:3343-3350(2015)

155 - A molecular network for functional versatility of HECATE transcription factors. Gaillochet C., Jamge S., van der Wal F., Angenent G., Immink R., Lohmann J.U. Plant J. 95:57-70(2018)

156 - The LNK gene family: at the crossroad between light signaling and the circadian clock. de Leone M.J., Hernando C.E., Romanowski A., García-Hourquet M., Careno D., Casal J., Rugnone M., Mora-García S., Yanovsky M.J. Genes 10:2(2019)

157 - Biosynthesis and emission of terpenoid volatiles from Arabidopsis flowers. Chen F., Tholl D., D'Auria J.C., Farooq A., Pichersky E., Gershenzon J. Plant Cell 15:481-494(2003)

158 - The Arabidopsis root stele transporter NPF2.3 contributes to nitrate translocation to shoots under salt stress. Taochy C., Gaillard I., Ipotesi E., Oomen R., Leonhardt N., Zimmermann S., Peltier J.B., Szponarski W., Simonneau T., Sentenac H., Gibrat R., Boyer J.C. Plant J. 83:466-479(2015)

159 - Dimerization properties of the transmembrane domains of Arabidopsis CRINKLY4 receptor-like kinase and homologs. Stokes K.D., Gururaj Rao A. Arch. Biochem. Biophys. 477:219-226(2008)

160 - Microarray expression profiling and functional characterization of AtTPS genes: duplicated *Arabidopsis thaliana* sesquiterpene synthase genes *At4g13280* and *At4g13300* encode root-specific and wound-inducible (Z)-gamma-bisabolene synthases. Ro D.K., Ehlting J., Keeling C.I., Lin R., Mattheus N., Bohlmann J. Arch. Biochem. Biophys. 448:104-116(2006)

161 - BBX28 negatively regulates photomorphogenesis by repressing HY5 activity and itself undergoes COP1-mediated degradation in Arabidopsis. Lin F., Jiang Y., Li J., Yan T., Fan L., Liang J.S., Chen Z.J., Xu D., Deng X.W. Plant Cell 30:2006-2019(2018)

162 - Antagonistic jacalin-related lectins regulate the size of ER body-type beta-glucosidase complexes in *Arabidopsis thaliana*. Nagano A.J., Fukao Y., Fujiwara M., Nishimura M., Hara-Nishimura I. Plant Cell Physiol. 49:969-980(2008)

163 - LNK genes integrate light and clock signaling networks at the core of the Arabidopsis oscillator. Rugnone M.L., Faigon Soverna A., Sanchez S.E., Schlaen R.G., Hernando C.E., Seymour D.K., Mancini E., Chernomoretz A., Weigel D., Mas P., Yanovsky M.J. Proc. Natl. Acad. Sci. U.S.A. 110:12120-12125(2013)

164 - AtCaM4 interacts with a Sec14-like protein, PATL1, to regulate freezing tolerance in Arabidopsis in a CBF-independent manner. Chu M., Li J., Zhang J., Shen S., Li C., Gao Y., Zhang S. J. Exp. Bot. 69:5241-5253(2018)

165 - Effects of COR6.6 and COR15am polypeptides encoded by COR (cold-regulated) genes of *Arabidopsis thaliana* on dehydration-induced phase transitions of phospholipid membranes. Webb M.S., Gilmour S.J., Thomashow M.F., Steponkus P.L. Plant Physiol. 111:301-312(1996)

166 - Cytochrome P450 CYP710A encodes the sterol C-22 desaturase in Arabidopsis and tomato. Morikawa T., Mizutani M., Aoki N., Watanabe B., Saga H., Saito S., Oikawa A., Suzuki H., Sakurai N., Shibata D., Wadano A., Sakata K., Ohta D. Plant Cell 18:1008-1022(2006)

167 - DREB1A/CBF3 is repressed by transgene-induced DNA methylation in the Arabidopsis *ice1-1* mutant. Kidokoro S., Kim J.S., Ishikawa T., Suzuki T., Shinozaki K., Yamaguchi-Shinozaki K. Plant Cell 32:1035-1048(2020)

168 - An Arabidopsis TIR-lectin two-domain protein confers defence properties against Tetranychus urticae. Santamaria M.E., Martinez M., Arnaiz A., Rioja C., Burow M., Grbic V., Diaz, I. Plant Physiol. 179:1298-1314(2019)

169 - Galactinol and raffinose constitute a novel function to protect plants from oxidative damage. Nishizawa A., Yabuta Y., Shigeoka S. Plant Physiol. 147:1251-1263(2008)

170 - Galactinol as marker for seed longevity. de Souza Vidigal D., Willems L., van Arkel J., Dekkers B.J.W., Hilhorst H.W.M., Bentsink L. Plant Sci. 246:112-118(2016)

171 - Natural variation in a polyamine transporter determines paraquat tolerance in Arabidopsis. Fujita M., Fujita Y., Iuchi S., Yamada K., Kobayashi Y., Urano K., Kobayashi M., Yamaguchi-Shinozaki K., Shinozaki K. Proc. Natl. Acad. Sci. U.S.A. 109:6343-6347(2012)

172 - PSEUDO-RESPONSE REGULATORS 9, 7, and 5 are transcriptional repressors in the Arabidopsis circadian clock. Nakamichi N., Kiba T., Henriques R., Mizuno T., Chua N.H., Sakakibara H. Plant Cell 22:594-605(2010)

173 - The Arabidopsis nitrate transporter NRT1.7, expressed in phloem, is responsible for source-to-sink remobilization of nitrate. Fan S.C., Lin C.S., Hsu P.K., Lin S.H., Tsay Y.F. Plant Cell 21:2750-2761(2009)

174 - Characterization of a root-specific Arabidopsis terpene synthase responsible for the formation of the volatile monoterpene 1,8-cineole. Chen F., Ro D.K., Petri J., Gershenzon J., Bohlmann J., Pichersky E., Tholl D. Plant Physiol. 135:1956-1966(2004)

175 - Susceptibility to Verticillium longisporum is linked to monoterpene production by TPS23/27 in Arabidopsis. Roos J., Bejai S., Mozūraitis R., Dixelius C. Plant J. 81:572-585(2015)

176 - The photosystem II repair cycle requires FtsH turnover through the EngA GTPase. Kato Y., Hyodo K., Sakamoto W. Plant Physiol. 178:596-611(2018)

177 - TSA1 interacts with CSN1/CSN and may be functionally involved in Arabidopsis seedling development in darkness. Li W., Zang B., Liu C., Lu L., Wei N., Cao K., Deng X.W., Wang X. J. Genet. Genomics 38:539-546(2011)

178 - Jasmonic acid-inducible TSA1 facilitates ER body formation. Geem K.R., Kim D.H., Lee D.W., Kwon Y., Lee J., Kim J.H., Hwang I. Plant J. 97:267-280(2019)

179 - Molecular cloning, phylogenetic analysis, expressional profiling and in vitro studies of TINY2 from Arabidopsis thaliana. Wei G., Pan Y., Lei J., Zhu Y.X. J. Biochem. Mol. Biol. 38:440-6(2005)

180 - Genetic and molecular identification of genes required for female gametophyte development and function in Arabidopsis. Pagnussat G.C., Yu H.J., Ngo Q.A., Rajani S., Mayalagu S., Johnson C.S., Capron A., Xie L.F., Ye D., Sundaresan V. Development 132:603-14(2005)

181 - Overexpression of the *TIR-X* gene results in a dwarf phenotype and activation of defense-related gene expression in Arabidopsis thaliana. Kato H., Saito T., Ito H., Komeda Y., Kato A.J. Plant Physiol. 171:382-388(2014)

182 - Polyamines in the life of Arabidopsis: profiling the expression of S-adenosylmethionine decarboxylase (SAMDC) gene family during its life cycle. Majumdar R., Shao L., Turlapati S.A., Minocha S.C. BMC Plant Biol. 17:264(2017)

183 - A putative hydroxysteroid dehydrogenase involved in regulating plant growth and development. Li F., Asami T., Wu X., Tsang E.W., Cutler A.J. Plant Physiol. 145:87-97(2007)

184 - A novel family of cys-rich membrane proteins mediates cadmium resistance in Arabidopsis. Song W.Y., Martinoia E., Lee J., Kim D., Kim D.Y., Vogt E., Shim D., Choi K.S., Hwang I., Lee Y. Plant Physiol. 135:1027-1039(2004)

185 - Regulation of HSD1 in seeds of *Arabidopsis thaliana*. Baud S., Dichow N.R., Kelemen Z., d'Andréa S., To A., Berger N., Canonge M., Kronenberger J., Viterbo D., Dubreucq B., Lepiniec L., Chardot T., Miquel M. Plant Cell Physiol. 50:1463-1478(2009)

186 - The Arabidopsis a4 subfamily of lectin receptor kinases negatively regulates abscisic acid response in seed germination. Xin Z., Wang A., Yang G., Gao P., Zheng Z.L. Plant Physiol. 149:434-444(2009)

187 - Arabidopsis ABCG28 is required for the apical accumulation of reactive oxygen species in growing pollen tubes. Do T.H.T., Choi H., Palmgren M., Martinoia E., Hwang J.U., Lee Y. Proc. Natl. Acad. Sci. U.S.A. 116:12540-12549(2019)

188 - A HECT E3 ubiquitin ligase negatively regulates Arabidopsis leaf senescence through degradation of the transcription factor WRKY53. Miao Y., Zentgraf U. Plant J. 63:179-188(2010)

189 - Arabidopsis WRKY53, a node of multi-layer regulation in the network of senescence. Zentgraf U., Doll J. Plants 8:578(2019)

190 - PETAL LOSS, a trihelix transcription factor gene, regulates perianth architecture in the Arabidopsis flower. Brewer P.B., Howles P.A., Dorian K., Griffith M.E., Ishida T., Kaplan-Levy R.N., Kilinc A., Smyth D.R. Development 131:4035-4045(2004)

191 - A gain-of-function mutation of transcriptional factor PTL results in curly leaves, dwarfism and male sterility by affecting auxin homeostasis. Li X., Qin G., Chen Z., Gu H., Qu L.J. Plant Mol. Biol. 66:315-327(2008)

192 - Transcriptional regulatory framework for vascular cambium development in Arabidopsis roots. Zhang J., Eswaran G., Alonso-Serra J., Kucukoglu M., Xiang J., Yang W., Elo A., Nieminen K., Damén T., Joung J.G., Yun J.Y., Lee J.H., Ragni L., Barbier de Reuille P., Ahnert S.E., Lee J.Y., Mähönen A.P., Helariutta Y. Nat. Plants 5:1033-1042(2019)

193 - The Arabidopsis sn-1-specific mitochondrial acylhydrolase AtDLAH is positively correlated with seed viability. Seo Y.S., Kim E.Y., Kim W.T. J. Exp. Bot. 62:5683-5698(2011)

194 - Overexpression of AtCSP4 affects late stages of embryo development in Arabidopsis. Yang Y., Karlson D.T. J. Exp. Bot. 62:2079-2091(2011)

195 - Storage protein accumulation in the absence of the vacuolar processing enzyme family of cysteine proteases. Gruis D., Schulze J., Jung R. Plant Cell 16:270-290(2004)

196 - The Arabidopsis ATR1 Myb transcription factor controls indolic glucosinolate homeostasis. Celenza J.L., Quiel J.A., Smolen G.A., Merrikh H., Silvestro A.R., Normanly J., Bender J. Plant Physiol. 137:253-262(2005)

197 – The role of MYB34, MYB51 and MYB122 in the regulation of camalexin biosynthesis in Arabidopsis thaliana. Frerigmann H., Glawischnig E., Gigolashvili T. Front. Plant Sci. 6:654 (2015)

198 - A genome-wide analysis of Arabidopsis Rop-interactive CRIB motif-containing proteins that act as Rop GTPase targets. Wu G., Gu Y., Li S., Yang Z. Plant Cell 13:2841-2856(2001)

199 - Glutathionylation inhibits catalytic activity of Arabidopsis β-amylase3 but not paralog β-amylase1. Storm A.R., Kohler M.R., Berndsen C.E., Monroe J.D. Biochemistry 57:711-721(2018)

200 - The HIRA complex that deposits the histone H3.3 is conserved in Arabidopsis and facilitates transcriptional dynamics. Nie X., Wang H., Li J., Holec S., Berger F. Biol. Open 3:794-802(2014)

201 - The redox-sensitive chloroplast trehalose-6-phosphate phosphatase AtTPPD regulates salt stress tolerance. Krasensky J., Broyart C., Rabanal F., Jonak C. Antioxid. Redox Signal. 21:1289-1304(2014)

202 - Two homologous INDOLE-3-ACETAMIDE (IAM) HYDROLASE genes are required for the auxin effects of IAM in Arabidopsis. Gao Y., Dai X., Aoi Y., Takebayashi Y., Yang L., Guo X., Zeng Q., Yu H., Kasahara H., Zhao Y. J. Genet. Genomics 47:157-165(2020)

203 - SCF E3 ligase PP2-B11 plays a positive role in response to salt stress in Arabidopsis. Jia F., Wang C., Huang J., Yang G., Wu C., Zheng C. J. Exp. Bot. 66:4683-4697(2015)

204 - Salt-induced stabilization of EIN3/EIL1 confers salinity tolerance by deterring ROS accumulation in Arabidopsis. Peng J., Li Z., Wen X., Li W., Shi H., Yang L., Zhu H., Guo H. PLoS Genet.10:e1004664(2014)

205 - The plant specific transcription factors CBP60g and SARD1 are targeted by a Verticillium secretory protein VdSCP41 to modulate immunity. Qin J., Wang K., Sun L., Xing H., Wang S., Li L., Chen S., Guo H.S., Zhang J. Elife 7:e34902(2018)

206 - The transcription factor HIG1/MYB51 regulates indolic glucosinolate biosynthesis in *Arabidopsis thaliana*. Gigolashvili T., Berger B., Mock H.P., Mueller C., Weisshaar B., Fluegge U.I. Plant J. 50:886-901(2007)

207 - Changing substrate specificity and iteration of amino acid chain elongation in glucosinolate biosynthesis through targeted mutagenesis of Arabidopsis methylthioalkylmalate synthase1. Petersen A., Hansen L.G., Mirza N., Crocoll C., Mirza O.A., Halkier B.A. Biosci. Rep. 39:0446(2019)

208 - The STRUCTURAL MAINTENANCE OF CHROMOSOMES 5/6 complex promotes sister chromatid alignment and homologous recombination after DNA damage in Arabidopsis thaliana. Watanabe K., Pacher M., Dukowic S., Schubert V., Puchta H., Schubert I. Plant Cell 21:2688-2699(2009)

209 - Biochemical and molecular characterization of flavonoid 7-sulfotransferase from *Arabidopsis thaliana*. Gidda S.K., Varin L. Plant Physiol. Biochem. 44:628-636(2006)

210 - ERF105 is a transcription factor gene of Arabidopsis thaliana required for freezing tolerance and cold acclimation. Bolt S., Zuther E., Zintl S., Hincha D.K., Schmulling T. Plant Cell Environ. 40:108-120(2017)

211 - ABI4 represses the expression of type-A ARRs to inhibit seed germination in Arabidopsis. Huang X., Zhang X., Gong Z., Yang S., Shi Y. Plant J. 89:354-365(2017)

212 - Proteasome-mediated turnover of Arabidopsis MED25 is coupled to the activation of FLOWERING LOCUS T transcription. Inigo S., Giraldez A.N., Chory J., Cerdan P.D. Plant Physiol. 160:1662-1673(2012)

213 - CYP707A3, a major ABA 8'-hydroxylase involved in dehydration and rehydration response in *Arabidopsis thaliana*. Umezawa T., Okamoto M., Kushiro T., Nambara E., Oono Y., Seki M., Kobayashi M., Koshiba T., Kamiya Y., Shinozaki K. Plant J. 46:171-182(2006)

214 - Glutathione conjugates in the vacuole are degraded by gamma-glutamyl transpeptidase GGT3 in Arabidopsis. Ohkama-Ohtsu N., Zhao P., Xiang C., Oliver D.J. Plant J. 49:878-888(2007)

215 - CENTRIN2 interacts with the Arabidopsis homolog of the human XPC protein (AtRAD4) and contributes to efficient synthesis-dependent repair of bulky DNA lesions. Liang L., Flury S., Kalck V., Hohn B., Molinier J. Plant Mol. Biol. 61:345-356(2006)

216 - A signal cascade originated from epidermis defines apical-basal patterning of Arabidopsis shoot apical meristems. Han H., Yan A., Li L., Zhu Y., Feng B., Liu X., Zhou Y. Nat. Commun. 11:1214(2020)

217 - Phosphorylation of Arabidopsis ubiquitin ligase ATL31 is critical for plant C/N-nutrient response and controls the stability of 14-3-3 proteins. Yasuda S., Sato T., Maekawa S., Aoyama S., Fukao Y., Yamaguchi J. J. Biol. Chem. 289:15179-15193(2014)

218 - AtRAB-H1b and AtRAB-H1c GTPases, homologues of the yeast Ypt6, target reporter proteins to the Golgi when expressed in Nicotiana tabacum and Arabidopsis thaliana. Johansen J.N., Chow C.M., Moore I., Hawes C. J. Exp. Bot. 60:3179-3193(2009)

219 - The AtGRXS3/4/5/7/8 glutaredoxin gene cluster on *Arabidopsis thaliana* chromosome 4 is coordinately regulated by nitrate and appears to control primary root. Walters L.A., Escobar M.A. Plant Signal. Behav. 11:e1171450(2016)

220 - Malonylation is a key reaction in the metabolism of xenobiotic phenolic glucosides in Arabidopsis and tobacco. Taguchi G., Ubukata T., Nozue H., Kobayashi Y., Takahi M., Yamamoto H., Hayashida N. Plant J. 63:1031-1041(2010)

221 - Characterization of an Arabidopsis enzyme family that conjugates amino acids to indole-3-acetic acid. Staswick P.E., Serban B., Rowe M., Tiryaki I., Maldonado M.T., Maldonado M.C., Suza W. Plant Cell 17:616-627(2005)

222 - Auxin controls Arabidopsis adventitious root initiation by regulating jasmonic acid homeostasis. Gutierrez L., Mongelard G., Floková K., Pacurar D.I., Novák O., Staswick P., Kowalczyk M., Pacurar M., Demailly H., Geiss G., Bellini C. Plant Cell 24:2515-2527(2012)

223 - Evolution of the small family of alternative splicing modulators nuclear speckle RNA-binding proteins in plants. Lucero L., Bazin J., Rodriguez Melo J., Ibanez F., Crespi M.D., Ariel F. Genes 11:207(2020)

224 - Cryptic variation in RNA-directed DNA-methylation controls lateral root development when auxin signalling is perturbed. Shahzad Z., Eaglesfield R., Carr C., Amtmann A. Nat. Commun. 11:218(2020)

225 - Nitrile-specifier proteins involved in glucosinolate hydrolysis in *Arabidopsis thaliana*. Kissen R., Bones A.M. J. Biol. Chem. 284:12057-12070(2009)

226 - PINOID-mediated signaling involves calcium-binding proteins. Benjamins R., Ampudia C.S., Hooykaas P.J., Offringa R. Plant Physiol. 132:1623-1630(2003)

227 - The plant WNK gene family and regulation of flowering time in Arabidopsis. Wang Y., Liu K., Liao H., Zhuang C., Ma H., Yan X. Plant Biol. 10:548-562(2008)

228 - Genomic and functional characterization of the oas gene family encoding O-acetylserine (thiol) lyases, enzymes catalyzing the final step in cysteine biosynthesis in *Arabidopsis thaliana*. Jost R., Berkowitz O., Wirtz M., Hopkins L., Hawkesford M.J., Hell R. Gene 253:237-247(2000)

229 - Interactions between the S-Domain receptor kinases and AtPUB-ARM E3 ubiquitin ligases suggest a conserved signaling pathway in Arabidopsis. Samuel M.A., Mudgil Y., Salt J.N., Delmas F., Ramachandran S., Chilelli A., Goring D.R. Plant Physiol. 147:2084-2095(2008)

230 - Phosphorylation of a WRKY transcription factor by two pathogen-responsive MAPKs drives phytoalexin biosynthesis in Arabidopsis. Mao G., Meng X., Liu Y., Zheng Z., Chen Z., Zhang S. Plant Cell 23:1639-1653(2011)

231 - Arabidopsis WRKY33 is a key transcriptional regulator of hormonal and metabolic responses toward Botrytis cinerea infection. Birkenbihl R.P., Diezel C., Somssich I.E. Plant Physiol. 159:266-285(2012)

232 - Functional dissection of the PROPEP2 and PROPEP3 promoters reveals the importance of WRKY factors in mediating microbe-associated molecular pattern-induced expression. Logemann E., Birkenbihl R.P., Rawat V., Schneeberger K., Schmelzer E., Somssich I.E. New Phytol. 198:1165-1177(2013)

233 - Phosphate homeostasis and root development in Arabidopsis are synchronized by the zinc finger transcription factor ZAT6. Devaiah B.N., Nagarajan V.K., Raghothama K.G. Plant Physiol. 145:147-159(2007)

234 - A signal cascade originated from epidermis defines apical-basal patterning of Arabidopsis shoot apical meristems. Han H., Yan A., Li L., Zhu Y., Feng B., Liu X., Zhou Y. Nat. Commun. 11:1214(2020)

235 - Arabidopsis ABCG transporters, which are required for export of diverse cuticular lipids, dimerize in different combinations. McFarlane H.E., Shin J.J., Bird D.A., Samuels A.L. Plant Cell 22:3066-3075(2010)

236 - A trihelix DNA binding protein counterbalances hypoxia-responsive transcriptional activation in Arabidopsis. Giuntoli B., Lee S.C., Licausi F., Kosmacz M., Oosumi T., van Dongen J.T., Bailey-Serres J., Perata P. PLoS Biol. 12:e100195(2014)

237 - D-glycerate 3-kinase, the last unknown enzyme in the photorespiratory cycle in Arabidopsis, belongs to a novel kinase family. Boldt R., Edner C., Kolukisaoglu U., Hagemann M., Weckwerth W., Wienkoop S., Morgenthal K., Bauwe H. Plant Cell 17:2413-2420(2005)

238 - Overview of OVATE FAMILY PROTEINS, a novel class of plant-specific growth regulators. Wang S., Chang Y., Ellis B. Front. Plant Sci. 7:417(2016)

239 - AXR1 promotes the Arabidopsis cytokinin response by facilitating ARR5 proteolysis. Li Y., Kurepa J., Smalle J. Plant J. 74:13-24(2013)

240 - Multifaceted role of cycling DOF factor 3 (CDF3) in the regulation of flowering time and abiotic stress responses in Arabidopsis. Corrales A.R., Carrillo L., Lasierra P., Nebauer S.G., Dominguez-Figueroa J., Renau-Morata B., Pollmann S., Granell A., Molina R.V., Vicente-Carbajosa J., Medina J. Plant Cell Environ. 40:748-764(2017)

241 - GLABROUS INFLORESCENCE STEMS3 (GIS3) regulates trichome initiation and development in Arabidopsis. Sun L., Zhang A., Zhou Z., Zhao Y., Yan A., Bao S., Yu H., Gan Y. New Phytol. 206:220-230(2015)

242 - The MYB96 transcription factor regulates cuticular wax biosynthesis under drought conditions in Arabidopsis. Seo P.J., Lee S.B., Suh M.C., Park M.J., Go Y.S., Park C.M. Plant Cell 23:1138-1152(2011)

243 - MYB96 recruits the HDA15 protein to suppress negative regulators of ABA signaling in Arabidopsis. Lee H.G., Seo P.J. Nat. Commun. 10:1713(2019)

244 - CYSTM, a novel non-secreted cysteine-rich peptide family, involved in environmental stresses in Arabidopsis thaliana. Xu Y., Yu Z., Zhang D., Huang J., Wu C., Yang G., Yan K., Zhang S., Zheng C. Plant Cell Physiol. 59:423-438(2018)

**C - more abundant transcripts in *dorn-1.1***


245 - Xyloglucan endotransglucosylase-hydrolase17 interacts with xyloglucan endotransglucosylase-hydrolase31 to confer xyloglucan endotransglucosylase action and affect aluminum sensitivity in Arabidopsis. Zhu X.F., Wan J.X., Sun Y., Shi Y.Z., Braam J., Li G.X., Zheng S.J. Plant Physiol. 165:1566-1574 (2014)

246 - Functionally redundant LNG3 and LNG4 genes regulate turgor-driven polar cell elongation through activation of XTH17 and XTH24. Lee Y.K., Rhee J.Y., Lee S.H., Chung G.C., Park S.J., Segami S., Maeshima M., Choi G. Plant Mol Biol 97:23-36 (2018)

247 - Perception of root-derived peptides by shoot LRR-RKs mediates systemic N- demand signaling. Tabata R., Sumida K., Yoshii T., Ohyama K., Shinohara H., Matsubayashi Y. Science 346:343-346 (2014)

248 - CEP5 and XIP1/CEPR1 regulate lateral root initiation in Arabidopsis. Roberts I., Smith S., Stes E., De Rybel B., Staes A., van de Cotte B., Njo M.F., Dedeyne L., Demol H., De Smet I. J. Exp. Bot. 67:4889-4899 (2016)

249 - CEP-CEPR1 signalling inhibits the sucrose-dependent enhancement of lateral root growth. Chapman K., Taleski M., Ogilvie H.A., Imin N., Djordjevic M.A. J Exp Bot 70:3955-3967 (2019)

250 - Agrobacterium-mediated root transformation is inhibited by mutation of an Arabidopsis cellulose synthase-like gene. Zhu Y., Nam J., Carpita N.C., Matthysse A.G., Gelvin S.B. Plant Physiol. 133:1000-1010 (2003)

251 - Arabidopsis mannan synthase CSLA9 and glucan synthase CSLC4 have opposite orientations in the Golgi membrane. Davis J., Brandizzi F., Liepman A.H., Keegstra K. Plant J 64:1028-1037 (2010)

252 - Cell wall glucomannan in Arabidopsis is synthesised by CSLA glycosyltransferases, and influences the progression of embryogenesis. Goubet F., Barton C.J., Mortimer J.C., Yu X., Zhang Z., Miles G.P., Richens J., Liepman A.H., Seffen K., Dupree P. Plant J 60:527-538 (2009)

253 - Genome-wide Association Study Reveals that the Aquaporin NIP1;1 Contributes to Variation in Hydrogen Peroxide Sensitivity in Arabidopsis thaliana. Sadhukhan A., Kobayashi Y., Nakano Y., Iuchi S., Kobayashi M., Sahoo L., Koyama H. Mol Plant 10:1082-1094 (2017)

254 - CYP83A1 and CYP83B1, two nonredundant cytochrome P450 enzymes metabolizing oximes in the biosynthesis of glucosinolates in Arabidopsis. Naur P., Petersen B.L., Mikkelsen M.D., Bak S., Rasmussen H., Olsen C.E., Halkier B.A. Plant Physiol. 133:63-72 (2003)

255 - Mitochondrial Pyruvate Carriers Prevent Cadmium Toxicity by Sustaining the TCA Cycle and Glutathione Synthesis. He L., Jing Y., Shen J., Li X., Liu H., Geng Z., Wang M., Li Y., Chen D., Zhang W. Plant Physiol 180:198-211 (2019)

256 - Arabidopsis Glutathione-S-Transferases GSTF11 and GSTU20 Function in Aliphatic Glucosinolate Biosynthesis. Zhang A., Luo R., Li J., Miao R., An H., Yan X., Pang Q. Front Plant Sci12:816233 (2022)

257 - Silent S-Type Anion Channel Subunit SLAH1 Gates SLAH3 Open for Chloride Root-to-Shoot Translocation. Cubero-Font P., Maierhofer T., Jaslan J., Rosales M.A., Espartero J., Diaz-Rueda P., Muller H.M., Hurter A.L., Al-Rasheid K.A., Geiger D. Curr Biol 26:2213-2220 (2016)

258 - Functional specification of Arabidopsis isopropylmalate isomerases in glucosinolate and leucine biosynthesis. He Y., Chen B., Pang Q., Strul J.M., Chen S. Plant Cell Physiol. 51:1480-1487 (2010)

259 - The plastidic bile acid transporter 5 is required for the biosynthesis of methionine-derived glucosinolates in *Arabidopsis thaliana*. Gigolashvili T., Yatusevich R., Rollwitz I., Humphry M., Gershenzon J., Fluegge U.-I. Plant Cell 21:1813-1829 (2009)

260 - Branched-chain aminotransferase4 is part of the chain elongation pathway in the biosynthesis of methionine-derived glucosinolates in Arabidopsis. Schuster J., Knill T., Reichelt M., Gershenzon J., Binder S. Plant Cell 18:2664-2679 (2006)

261 - A novel polyamine acyltransferase responsible for the accumulation of spermidine conjugates in Arabidopsis seed. Luo J., Fuell C., Parr A., Hill L., Bailey P., Elliott K., Fairhurst S.A., Martin C., Michael A.J. Plant Cell 21:318-333 (2009)

D - **less abundant transcripts in *dorn-1.1***

262 - EXPANSINA17 up-regulated by LBD18/ASL20 promotes lateral root formation during the auxin response. Lee H.W., Kim J. Plant Cell Physiol. 54:1600-1611(2013)

263 - PRX2 and PRX25, peroxidases regulated by COG1, are involved in seed longevity in Arabidopsis. Renard J., Martinez-Almonacid I., Sonntag A., Molina I., Moya-Cuevas J., Bissoli G., Munoz-Bertomeu J., Faus I., Ninoles R., Bueso E. Plant Cell Environ 43:315-326 (2020)

264 - Simultaneously disrupting *AtPrx2*, *AtPrx25* and *AtPrx71* alters lignin content and structure in Arabidopsis stem. Shigeto J., Itoh Y., Hirao S., Ohira K., Fujita K., Tsutsumi Y. J Integr Plant Biol 57:349-356 (2015)

265 - Disruption of AtOCT1, an organic cation transporter gene, affects root development and carnitine-related responses in Arabidopsis. Lelandais-Briere C., Jovanovic M., Torres G.A.M., Perrin Y., Lemoine R., Corre-Menguy F., Hartmann C. Plant J. 51:154-164 (2007)

266 - The putative high-affinity nitrate transporter NRT2.1 represses lateral root initiation in response to nutritional cues. Little D.Y., Rao H., Oliva S., Daniel-Vedele F., Krapp A., Malamy J.E. Proc. Natl. Acad. Sci. U.S.A. 102:13693-13698 (2005)

267 - A central role for the nitrate transporter NRT2.1 in the integrated morphological and physiological responses of the root system to nitrogen limitation in Arabidopsis. Remans T., Nacry P., Pervent M., Girin T., Tillard P., Lepetit M., Gojon A. Plant Physiol. 140:909-921 (2006)

268 - The homeobox genes ATHB12 and ATHB7 encode potential regulators of growth in response to water deficit in Arabidopsis. Olsson A.S.B., Engstroem P., Seoderman E. Plant Mol. Biol. 55:663-677 (2004)

269 - Identification of marneral synthase, which is critical for growth and development in Arabidopsis. Go Y.S., Lee S.B., Kim H.J., Kim J., Park H.Y., Kim J.K., Shibata K., Yokota T., Ohyama K., Suh M.C. Plant J 72:791-804 (2012)

270 - Linker histone variant HIS1-3 and WRKY1 oppositely regulate salt stress tolerance in Arabidopsis. Wu X., Xu J., Meng X., Fang X., Xia M., Zhang J., Cao S., Fan T. Plant Physiol. 189:1833-1847 (2022)

271 - Tonoplast intrinsic proteins AtTIP2;1 and AtTIP2;3 facilitate NH3 transport into the vacuole. Loque D., Ludewig U., Yuan L., von Wiren N. Plant Physiol 137:671-680 (2005)

272 - AtRD22 and AtUSPL1, members of the plant-specific BURP domain family involved in Arabidopsis thaliana drought tolerance. Harshavardhan V.T., Van Son L., Seiler C., Junker A., Weigelt-Fischer K., Klukas C., Altmann T., Sreenivasulu N., Baeumlein H., Kuhlmann M. PLoS ONE 9:E110065-E110065 (2014)

273 - Disruption of the Sugar Transporters AtSWEET11 and AtSWEET12 Affects Vascular Development and Freezing Tolerance in Arabidopsis. Le Hir R., Spinner L., Klemens P.A., Chakraborti D., de Marco F., Vilaine F., Wolff N., Lemoine R., Porcheron B., Bellini C. Mol Plant 8:1687-1690 (2015)

274 - The Arabidopsis stearoyl-acyl carrier protein-desaturase family and the contribution of leaf isoforms to oleic acid synthesis. Kachroo A., Shanklin J., Whittle E., Lapchyk L., Hildebrand D., Kachroo P. Plant Mol Biol 63:257-271 (2007)

275 - ABSCISIC ACID-DEFICIENT4 has an essential function in both cis-violaxanthin and cis-neoxanthin synthesis. Perreau F., Frey A., Effroy-Cuzzi D., Savane P., Berger A., Gissot L., Marion-Poll A. Plant Physiol 184:1303-1316 (2020)
