## Supplementary Figure S1 for "Receptor kinase LecRK-I.9 regulates cell wall remodelling and signalling during lateral root formation in *Arabidopsis*"

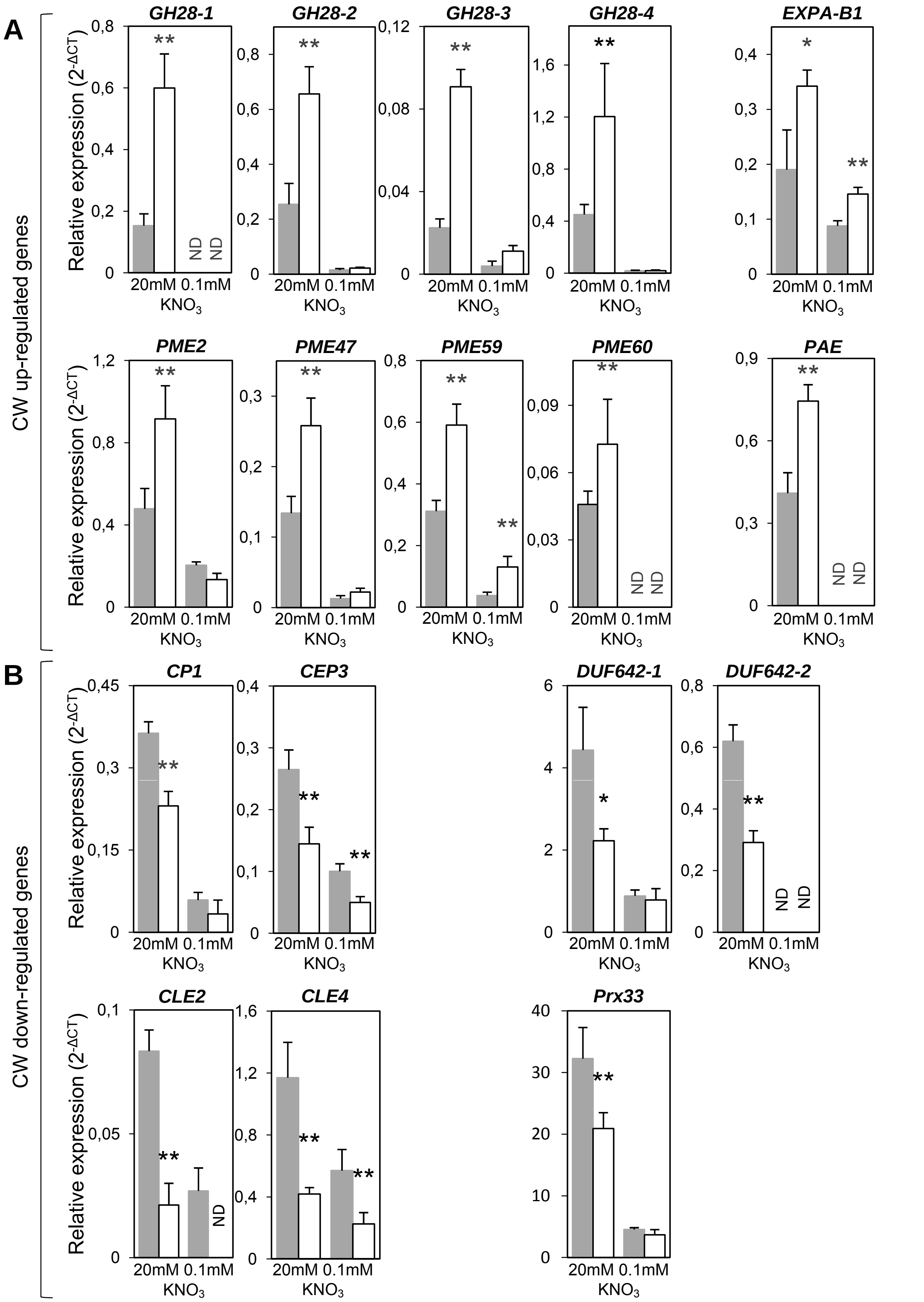


**Supplementary Fig. 1:** LecRK-I.9 regulates different sets of genes encoding cell wall proteins and peptides in roots. Transcript abundance was determined by RT-qPCR with cDNA generated from the roots of 7d-old seedlings. The target genes were selected from the transcriptomics analysis (Supplementary Dataset 1) devoted to the comparison between WT (grey bars) and *lecrk-I.9-1* (white bars): (A) up-regulated genes. (B) down-regulated genes. The means ± standard deviation of five biological replicates are shown. Asterisk denotes statistically significant differences according to Student's t-test (*$P$ < 0.05; **$P$ < 0.01) between *lecrk-I.9-1 versus* WT. ND: not detected. CW: cell wall. *GH28-1*: *At1g05650*. *GH28-2*: *At1g05660*. *GH28-3*: *At2g43870*. *GH28-4*: At2g43880. *DUF642-1*: *At2g34510*. *DUF642-2*: *At2g41810*.
