## Supplementary Figure S2 for "Receptor kinase LecRK-I.9 regulates cell wall remodelling and signalling during lateral root formation in *Arabidopsis*"

**A**

| Mean (%Dry mass) | | | | | | | | | | |
|---|---|---|---|---|---|---|---|---|---|---|
| **Genotype** | Rha | Fuc | Ara | Xyl | Man | Gal | total-Glc | resistant-Glc | UA | TS |
| **WT** | 0.5 | 0.4 | 3.9 | 2.5 | 0.8 | 2.0 | 11.5 | 9.6 | 6.0 | 27.6 |
| ***lecRK-I.9-1*** | 0.6 | 0.3 | 4.1 | 2.8 | 0.9 | 2.5 | 14.0 * | 12.4 * | 6.9 | 32.2 |
| ***lecRK-I.9-2*** | 0.5 | 0.4 | 3.9 | 2.7 | 0.9 | 2.1 | 12.2 | 9.9 | 6.5 | 29.2 |
| ***dnm-1*** | 0.5 | 0.4 | 3.6 | 2.4 | 0.7 | 2.1 | 11.0 | 9.5 | 7.5 | 28.2 |
| ***dnm-2*** | 0.5 | 0.3 | 3.3 | 2.4 | 0.9 | 2.1 | 11.9 | 9.9 | 7.3 | 28.8 |
| ***Ox-1*** | 0.5 | 0.4 | 3.8 | 2.5 | 0.9 | 2.3 | 11.2 | 9.4 | 6.6 | 28.2 |
| ***Ox-2*** | 0.4 | 0.4 | 3.4 | 2.2 | 0.7 | 2.0 | 10.3 | 8.3 | 6.7 | 26.2 |

**B**

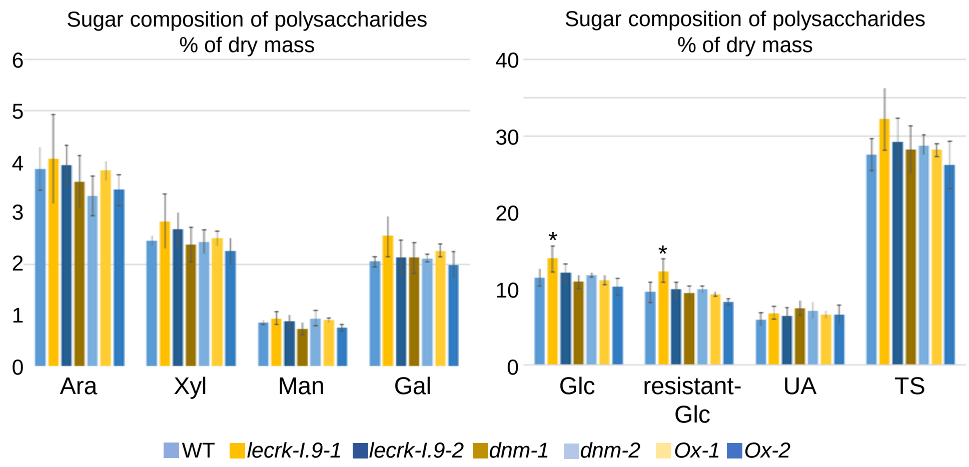


**Supplementary Fig. S2:** Monosaccharide contents of polysaccharides extracted from root tissues of different *A. thaliana* genotypes with modified *LecRK-I.9* level of expression. (A) The means of three biological replicates expressed as a percentage of dry mass are shown. Ara: arabinose; Fuc: fucose; Gal: galactose; Glc: glucose; Man: mannose; Rha: rhamnose; TS: total sugars; UA: uronic acids; Xyl: xylose. (B) The mean ± standard deviation of three biological replicates expressed as a percentage of dry mass are shown. Asterisk denotes statistically significant differences according to Student's t-test ($P < 0.05$) between *lecrk-I.9-1* plants *versus dnm-1* or *Ox-2* plants for total-Glc. and between *lecrk-I.9-1* plants *versus dnm-1. Ox-1* or *Ox-2* plants for resistant-Glc.
