## Supplementary Figure S3 for "Receptor kinase LecRK-I.9 regulates cell wall remodelling and signalling during lateral root formation in *Arabidopsis*"

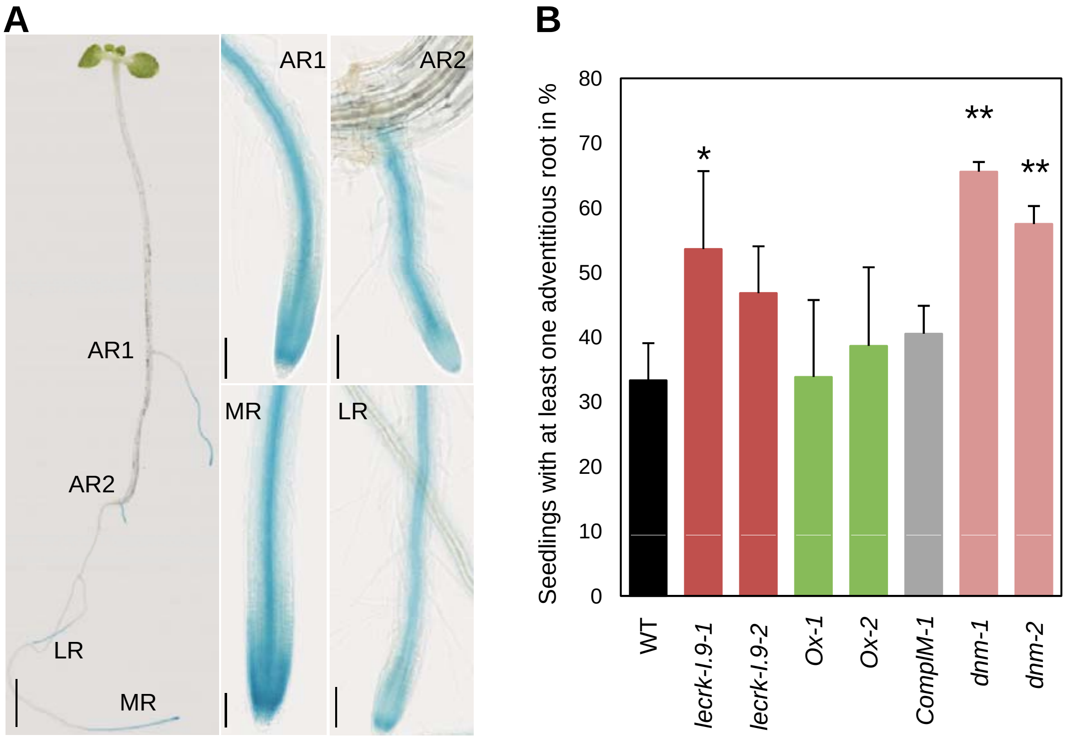


**Supplementary Fig. S3:** Adventitious root *lecrk-I.9* phenotype. (A) Representative seedlings that have developed adventitious roots. Close-up views show root apices of adventitious (AR1, AR2), lateral (LR), and main (MR) roots. The seedling was that of a *pLecRK-I.9:GUS* reporter line. The GUS enzymatic reaction time was 10 min. (B) Adventitious roots were counted in seedlings that were first etiolated in the dark for 4 days, and then transferred to the light for 7 days. The graph shows the percentage of seedlings of WT, knockout mutants (*lecrk-I.*9-1; *lecrk-I.*9-2), over-expressors (*Ox-1*; *Ox-2*), complemented (*ComplM-1*) and dominant negative mutants (*dnm-1*; *dnm-2*) lines having at least one adventitious root. The mean ± standard deviation of 65<n<122 seedlings is shown. Asterisks denote statistically significant differences according to Student's t-test (***P* < 0.01; **P*<0.05*)* between mutant, over-expressors, complemented and dominant negative mutant lines *vs* WT plants.
