## Supplementary Figure S4 for "Receptor kinase LecRK-I.9 regulates cell wall remodelling and signalling during lateral root formation in *Arabidopsis*"

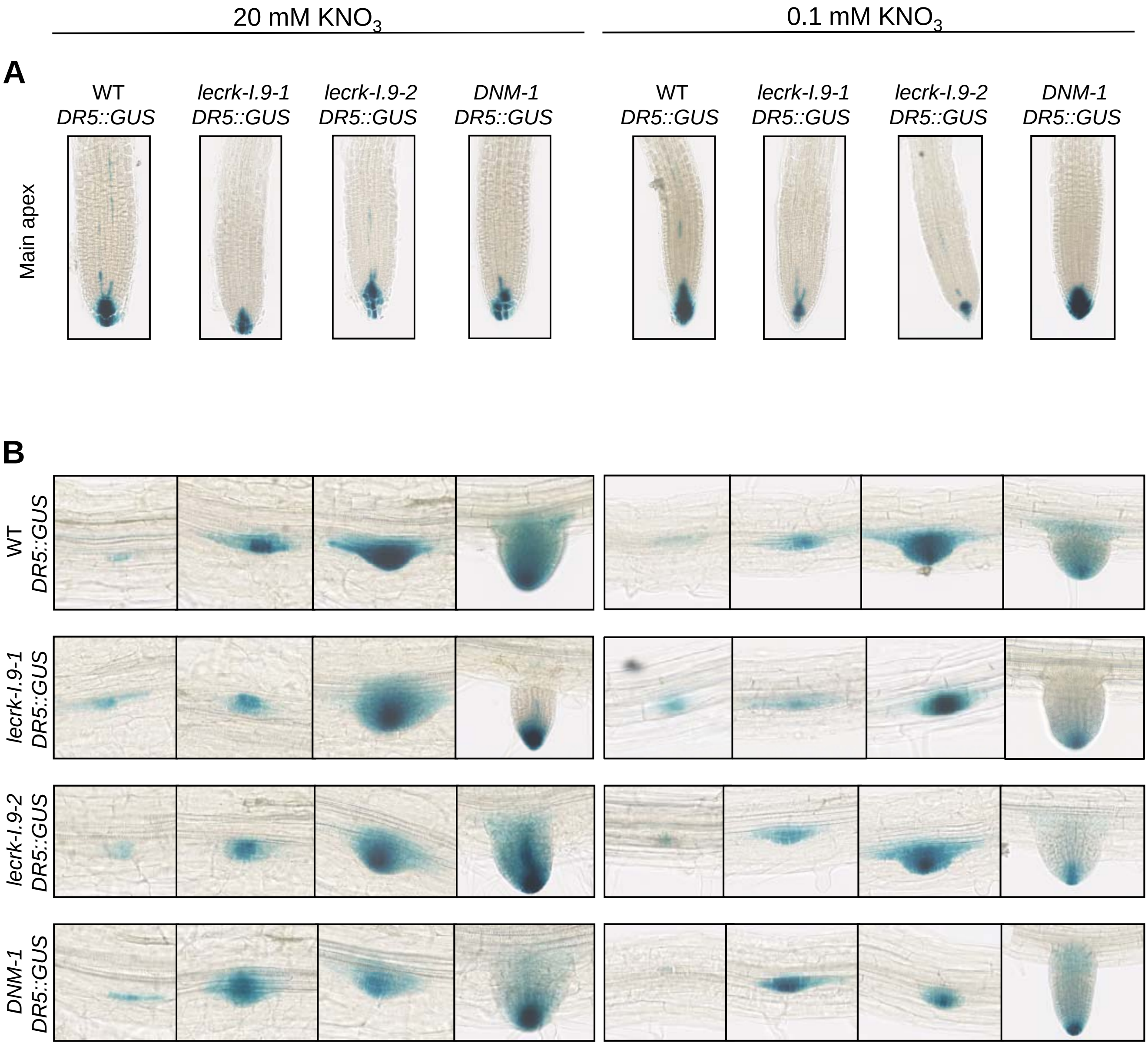


**Supplementary Fig. S4:** Histochemical staining of GUS activity in seedlings of different genotypes transformed with the auxin-responsive reporter *DR5::GUS.* (A) The GUS activity was detected in the apex of the main roots. (B) The GUS activity was also detected at different stages of lateral root development in seedlings of WT, knockout mutants (*lecrk-I.9-1*; *lecrk-I.9-2*) and dominant negative mutant (*DNM-1*) lines grown either on half Murashige and Skoog culture medium (20 mM $KNO_3$) or under nitrate deficiency (0.1 mM $KNO_3$). For each line, at least 30 seedlings were analysed.
